# Transcriptome-based lead generation, ligand- and structure-based prioritization and experimental validation of TLR5-activating molecules

**DOI:** 10.64898/2026.02.25.707690

**Authors:** Anuja Jain, Hungyo Hungharla, Naidu Subbarao, Vibha Tandon, Shandar Ahmad

## Abstract

Current *in silico* drug discovery protocols ubiquitously depend on lead generation using a ligand-based approach in which novel leads are generated by fragment-signature matching or by a structure-based search involving molecular docking and conformational dynamics. None of them incorporates cellular contexts in which these drugs ultimately operate, leaving the task to a later stage of optimization leading to a high failure rate. Incorporating systems-level responses of drugs in an early stage of lead generation can significantly address this concern but has not been sufficiently explored. In this work, we employ a systems-level approach using connectivity map (CMAP) library to generate leads against a challenging system of a TLR pathway. Starting with gene expression data of TLR5 activation by its natural ligand, we generated molecular leads using CMAP and rigorously analyzed their validity using ligand and structure-based approaches, and helping to prioritize top hits. Experimental validation using ELISA-based antibody assay confirmed the activation of TLR5 by *each of the top nine prioritized leads* with their dose-dependent patterns suggesting that some of them may actually interact with the TLR signaling pathway in a complex manner. Although, demonstrated on TLR5, the proposed framework is intuitively scalable to other lead generation and optimization tasks.

## 1 INTRODUCTION

Drug discovery process for known diseases follows a more or less well-established procedure that starts with the identification of a suitable drug target followed by high throughput screening, hit identification, lead optimization and finally the selection of a candidate or lead molecule for in vitro, in vivo and clinical trials **[Mohs and Greig, 2017]**. Many drug candidates fail clinical trials of safety and efficacy at a later stage of this protocol primarily because the initial and mid-stage screening do not account for the complex cellular conditions and their biological context **[Hughes et al., 2011]**. Introduction of cellular perturbations by a candidate drug at an early stage of leads can therefore reduce costs associated with drug discovery **[Hughes et al., 2011; mohs and Greig, 2017]**. In this context, the development of a high throughput library of expected transcriptome changes by drug candidates is a promising avenue for drug discovery but appears to be under-utilized **[Verbist et al., 2015; Seligmann 2003]**. Systems-level responses for hit molecules in this procedure involve comparison of gene expression profiles, which implicitly considers activation of signaling pathways and tissue or organ physiology at the outset of lead generation. This strategy of looking up a drug that is expected to mimic or reverse the transcriptome changes brought by Connectivity Map (CMap) **[Lamb et al., 2006]**, can be termed as systems-level or transcriptome-based drug discovery (TBDD) in contrast to the traditional approach of structure based drug discovery (SBDD). In a practical scenario TBDD can still be followed by SBDD or vice versa to optimize early stage lead generation process.

Apart from implicitly modeling systems-level context into early stage lead generation, a huge advantage of TBDD is also that it completely does away with the need for selecting a specific drug target; leave alone the availability of its three-dimensional structure **[Iorio et al., 2010; Dopazo 2014; Pabon et al., 2018]**. Thus TBDD is specially suited for targeting complex biological targets together and those with unknown structures or knowledge of mechanism of interactions **[Paananen and Fortino, 2020]**. When available, the multiple modalities such as structure, ligand template and TBDD connectivity will be a great advantage as high confidence leads, satisfying multiple criterion can be selected at an early stage.

Greatest challenge to TBDD and its integration with structural and ligand information is that an exact mechanism of the selected leads in modulating the expression of target genes is not always obvious. Only a little work has been done towards TBDD optimization and integration with SBDD **[Hughes et al., 2000; Waring et al., 2001]**.

As a test case for this work, we have attempted to discover potential drug candidates for a complex immunological target of Toll-like receptor (TLR5) and evaluated the validity of prioritized targets experimentally. Toll like receptors (TLRs) are a group of molecules, considered some of the most difficult and yet very promising groups of targets in infectious diseases and cancer immunotherapy **[O’Neill et al., 2009; Braunstein et al., 2018]**. They serve as pattern recognition receptors (PRRs) for pathogen-associated molecular patterns (PAMPs) whose stimulation leads to adaptive immune responses mediated by cytokines production, type I interferon and other mediators **[Akira et al., 2006]**. There are 10 members in human TLR family-named by their numeric suffixes i.e. TLR1 to TLR10 that are localized to the cell surface (TLR1, TLR2, TLR4, TLR5, and TLR6) or to intracellular compartments such as the endoplasmic reticulum, endosome, lysosome, or endolysosome (TLR3, TLR7, TLR8, and TLR9) **[Kumar et al., 2009; Moresco et al., 2011]**. Overall, TLR5 is a complex drug target system, whose potential is not fully exploited because of a complex interplay between different TLRs on the one hand and regulation of TLR5 expression by several adapters and members of signaling pathways on the other. This makes the SBDD approach difficult to apply on this system. We try to evaluate and establish TBDD approach to this difficult system and identify hits which can simplify the process of drug discovery against TLR5.

We specially considered TLR5 out of all the TLRs because it is expressed in immune as well as in non immune cells and useful structural and transcriptomic information is available in literature, which will be handy when some of the candidates are considered for clinical applications in the future. TLR5 expresses in immune cells such as mucosal macrophages and dendritic cells as well as non immune cells such as intestinal epithelial cells **[Monie, 2017; Yang and Yan, 2017]**. TLR5 activation has been intensely investigated in the context of radiation-induced toxicity, resulting in the development of the optimized flagellin derivative from *Salmonella enterica*, CBLB502 (CBLB) or Entolimod [**Burdelya LG et al., 2008; Melin et al., 2021**]. It is a polypeptide drug and has the ability to activate NF-κB while inhibiting the pro-apoptotic p53 pathway [**Hennessy et al., 2010**]. The treatment of CBLB502 leads to the expression of the strong natural antioxidant superoxide dismutase (SOD)2 and induces the expression of several radioprotective cytokines, such as GCSF, IL-6, and TNF-α [**Burdelya LG et al., 2008; Bai et al., 2019**]. Importantly, it does not disrupt the anti-tumor impact of radiation. Based on its immunostimulatory effect and protective properties, the use of CBLB502 has advanced towards clinical trials to promote antitumor responses, to reduce radiotherapy and/or chemotherapy-induced side effects [**Melin et al., 2021**].

Apart from the immunological and transcriptomic data, the X-ray crystal structure of full length human TLR5 is still not yet available in Protein Data Bank (PDB), geometry minimized model of (PDB ID: 3J0A) based on sequence homology fitted Electron Microscopy (EM) single particle reconstruction at 26 Å resolution is available. It is reported to be an asymmetric heterodimer (chain A and B) model with 22-858 residues, including 23-629 residues in ectodomain, 640-664 residues in transmembrane helical domain for both the chains and 681-845 residues in TIR domain for chain A only. This model includes two disulfide bonds for each chain at residues 583–610 and 585–629 **[Zhou et al., 2012]**. After this study was started, AlphaFold based protein structure database and tool developed by DeepMind and EMBL-EBI have become available (https://alphafold.ebi.ac.uk/). AlphaFold database and AI tool contains over 200 million protein structures, providing broad coverage of UniProt sequences belongs to 47 proteomes and allows open access to predict the 3D structure of protein from its amino acid sequence [**Jumper et al., 2021; Varadi et al., 2022**].

Primary approach of TBDD and its integration with traditional methods in this work refers to the use of CMap library, which currently consists of over 1.5M gene expression profiles from ∼5,000 small-molecule compounds, and ∼3,000 genetic reagents, rigorously tested in multiple cell types **[Zhang et al., 2022]**. Overall experimental design for this study is shown in Figure 1(A), and a general CMAP protocol is explained in Figure 1(B). CMap provides a method to compare gene expression (transcriptome) signatures of disease conditions with drug signatures in CMap library and predict the most suitable hit which can mimic or reverse the biological perturbation of interest. In its basic form, CMap method employs a non-parametric rank-ordered Kalmogorov-Smirnov statistical for this search **[Hollander and Wolfe 1999]**. A pattern-matching tool (CMap tool: http://clue.io/query) is also provided by CMap to detect these connection between diseases and therapeutic molecules of interest **[Lamb et al., 2006; Subramanian et al., 2017]**. CMap tool can be used to make two types of queries in respect to providing the lead compounds or perturbagens of similar as well as reverse gene expression profiles: (1) query of up-regulated genes (upG) and down-regulated genes (downG), called cmap1 method and (2) query of only upG, called cmap2 method in this work.

**Figure 1:**
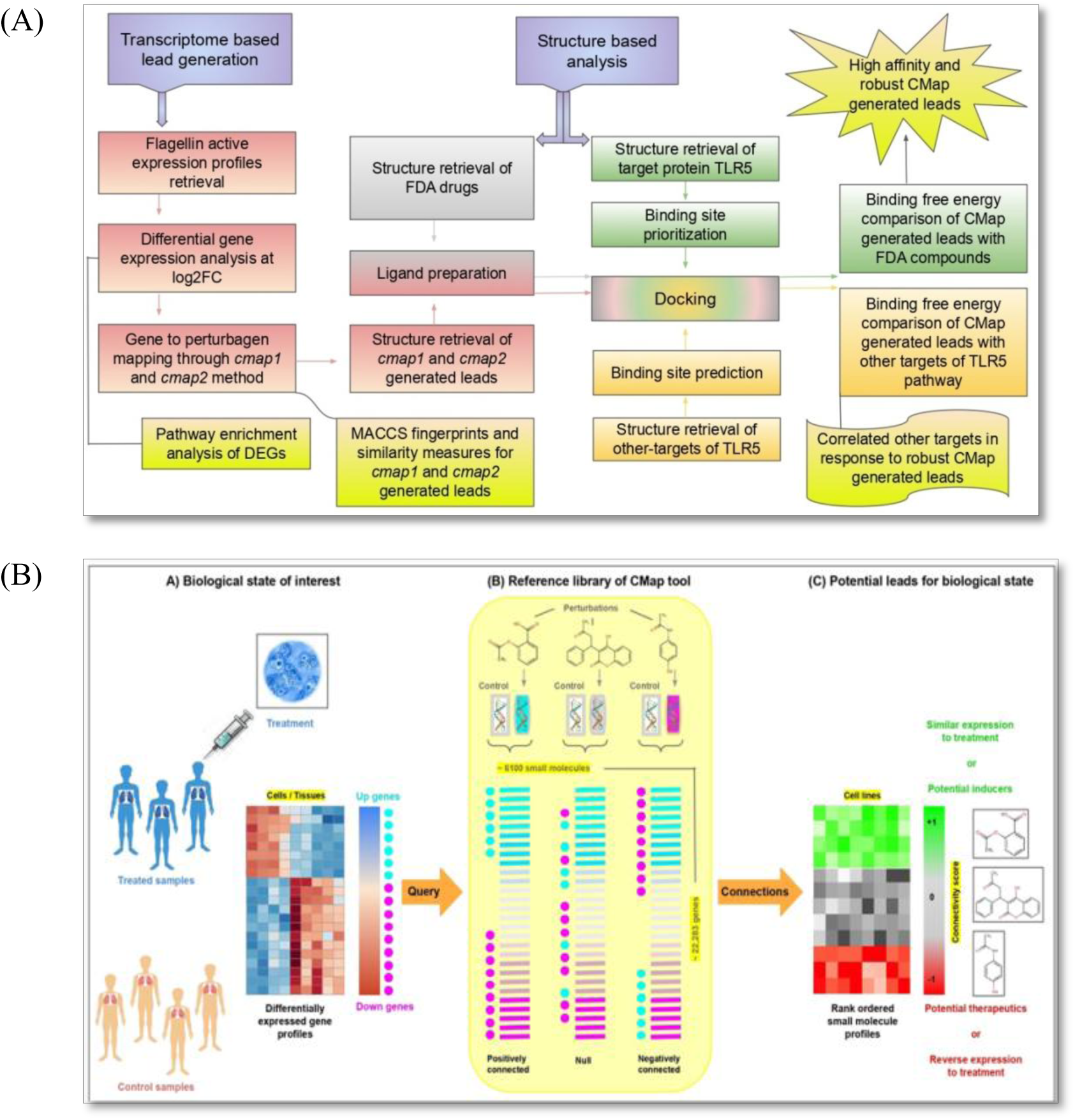
Overall lead generation protocol in this work. (A) Integrated design starting from Transcriptome-based lead generation, optimized by structure analysis (b) Detailed protocol for lead generation by Transcriptome based drug discovery (TBDD) (first column in (A)**. Overall** the method starts with a set of differentially expressed genes (DEGs) in a disease considering (upregulated and downregulated set; cmap1 method) or one direction of change (only the upregulated ones; cmap2 method). CMap tool then generates leads by mapoing these profiles to a pre-existing set of small molecules library. Leads generated by CMAP are then validated and optimized using standard structure-based approaches as outlined in (A)

As stated above, lead discovery using a CMAP based method for TLR5 was likely to overcome some of the limitations faced in SBDD-based drugs design **[Cheng et al., 2014; Musa et al., 2018]**. We implemented our strategy by determining lead molecules for TLR5 by using TBDD approach validated and prioritized using fragment enrichment in identified hits and molecular docking based SBDD. Prioritized hits were tested to activate TLR5 in cellular condition using and ELISA test and confirmed that all the nine prioritized hits using our integrated approach activated TLR5 in Calu cell lines. However, dose dependent pattern of activation was different in some cases suggesting that these drugs also interact with molecules other than TLR5 and thereby regulate this pathway in an indirect manner. An attempt to study molecular interactions with additional members of TLR5 pathway supported the presence of interactions between these hits and additional non-TLR5 proteins. However a more rigorous understanding for some of these leads activating TLR5 need to be understood in the future.

## 2. METHODS

Overall lead generation and optimization methodology adopted in this study is summarized in **Figure 1**. As shown, we aim to first generate leads using transcriptome based approach. To do so, we first identify the genes differentially expressed (DE) upon TLR5 activation by its natural ligand flagellin and using the selected DE genes, query CMAP database to generate leads which could mimic flagellin activation. As a parallel independent step, we carry out traditional structure-based screening. In this step, we start with molecular docking experiments on TLR5 directly using compound library. Finally, we combine the outcomes of transcriptome and structure based screening approaches and juxtaposed them in terms of their connectivity and docking scores. The objective here is to assess if CMAP method produces structurally compatible leads and then integrate the two outcomes to optimize CMAP leads. Individual steps involved in each of these are described in the following.

### 2.1 Transcriptome based lead generation

In this step (first column of Figure 1A, further explained in Figure 1B), gene expression profiles of typical TLR5 activation by flagellin have been used as a reference system. To ensure the quality of selected gene set, a pathway analysis was carried out on the DEGs identified in this step and subsequently, DEGs were used to query the connectivity map (CMAP) database to identify the leads. This process was achieved in the following manner:

#### 2.1.1 Dataset of flagellin active profiles

CMAP method depends on the choice of data set for selecting DEGs. Flagellin activation has been studied in various cell types, but it has been suggested that lung has the strongest host responses upon flagellin exposure [**Ishii and Akira 2008; Morris et al., 2009**]. Therefore, we started with the GE data of flagellin-activated Calu-3 cell lines from Gene Expression Omnibus (GEO). Calu-3 is a submucosal gland cell line, which was generated from a bronchial adenocarcinoma. There are two data series namely GSE133495 and GSE923 in GEO (https://www.ncbi.nlm.nih.gov/geo/) database which contain gene expression profiles of TLR5 activation through flagellin herein called flagellin active profiles. Both these experiments have been performed with legacy microarray technology. GSE133495, provides a comparison of the transcriptome profiles in Calu-3 cell lines in response to *pseudomonas aeruginosa* virulence factor flagellin that causes cystic fibrosis, expression data collected on Illumina HiSeq 2500. Only four samples with two samples belonging to control cell line (Ctrl_NS) and two stimulated with flagellin (Ctrl_Flag) are available from this study. On the other hand GSE923 contains five groups of samples namely Control, FRD1, FRD1234, FRD440 and FRD875, whose expression is reported on Affymetrix U133A arrays with infection modeled on *pseudomonas aeruginosa* strains. FRD440 group samples in this study were used to see the flagellin responses. This series contains a total of eight samples (4 control + 4 flagellin treated). Due to the challenges faced in multi-platform integration of microarray data and quantification of expression values, only the latter of the two with these eight samples (GSE923) was selected for subsequent analysis in this work.

#### 2.1.2 Identification of Differentially expressed genes (DEGs)

Expression profiles available under the above-mentioned source are already preprocessed and normalized and have not been further preprocessed for this work. In the original study, only the absent and present calls were reported for all the probes which have p-value <= 0.01 and used to identify DEGs. However, we assumed that for the current study, actual fold values of gene expression would be more suitable to create a comprehensive signature of flagellin which targets TLR5. In order to determine the DEGs afresh, we first assigned gene names to all the probe-level data using GSE923 annotation labels and the probes corresponding the highest GE value was selected as representative of the gene. Fold change was calculated between the average values of gene expression of each group (4 samples each). Finally, we collected DEGs (upG and downG) at log2 fold change (log2FC) threshold of 2 and plotted it using Srplot (https://www.bioinformatics.com.cn/plot_basic_scatter_plot_with_fc_lines_044_en).

#### 2.1.3 Pathway Enrichment analysis of DEGs

To ensure that the selected genes represent biological plausible perturbations, we looked at the enrichment of Kyoto Encyclopedia of Genes and Genomes (KEGG) pathways using “GO pathway enrichment analysis” module of SRplot, freely available from (https://www.bioinformatics.com.cn/basic_local_go_pathway_enrichment_analysis_122_en).

The “compareCluster” function of SRplot was used to identify the enriched pathways for each gene clusters with the strict filtering critereon cutoffs (p-value < 0.01 and FDR < 0.05).

#### 2.1.4 Finding CMap-generated leads using DEGs

CMAP queries were made using (a) only the upregulated DEGs from the above or (b) by pooling together the upregulated and downregulated DEGs. The two methods have been named as cmap1 and cmap2 respectively. So-called designated connectivity scores (CMap scores) were collected using CMAP interface (https://clue.io/**<u>, [Lamb et al., 2006]</u>**reference needed). CMap score is an internally normalized score having values from −1 to +1 that reflects the closeness or connection between the query GE profiles and a candidate lead. A particular lead compound from CMAP can either mimic (positive score) or reverse (negative score) the expression pattern of the biological state of interest. In our case, we selected top-scoring small molecules (positive score mimics) at a threshold of 0.9 on CMap score, ranked them by sorting and named them as “CMAP-generated leads”.

### 2.2 Chemical signature enrichment in transcriptome derived leads

In order to investigate the implicit structural signatures- if any- in CMAP hits, we examined the kind of structural patterns enriched in them. Purpose of this step was to assess if the CMAP generated compounds exhibit some pattern in terms of their chemical signatures, thereby addressing the concordance level between the two approaches. For this, we first computed similarity measures and enriched features among all the CMap-generated leads each of the lead sets resulting from cmap1 and cmap2 procedures, explained above. In order to assess the enriched structural patterns in these leads, we used the following steps.

#### 2.2.1 Structure retrieval of CMap-generated leads

All the 2D structures of CMap-generated leads were collected in Simplified Molecular Input Line Entry Specification (SMILE) format from the CMap (https://clue.io/command) library. Then we used the open source software OpenBabel (http://www.cheminfo.org/Chemistry/Cheminformatics/FormatConverter/index.html) to convert them into 3D Structural Data Format (SDF) and then identified the fragment contents of each of them using MACCS fingerprints.

#### 2.2.2 MACCS Fingerprint similarity calculation

Molecular ACCess System (MACCS) fingerprints are widely used to cluster drug candidates [**Gortari et al., 2017**]. There are two versions of these signatures based on 166 and 881 bit representations respectively. In our case, we have relatively smaller set of CMap-generated leads from both the methods (cmap1 and cmap2) therefore 166 bits MACCS keys fingerprints were found to be sufficient for the current analysis [**Gortari et al., 2017**]. To compare compounds represented by their fingerprints, Tanimoto coefficient was used as a similarity measures leading to a similarity matrix leads. [**Backman et al., 2011**]. “Chem”, “MACCSkeys” and “DataStructs” modules of python library RDKit (https://www.rdkit.org/docs/GettingStartedInPython.html) were used for computing fingerprint and similarity indices.

#### 2.2.3. Clustering of CMap-generated leads

Hierarchical clustering of compounds in fingerprints space was carried out by applying a ‘Ward linkage’ method using scipy.cluster.hierarchy library of python [**Hernández-Hernández and Ballester 2023**]. We cut the binary merged tree resulting from clustering at the distance of 5 and 15 to get robust and non-overlapping 166 bits MACCS fingerprints of cmap1 and cmap2 generated leads.

#### 2.2.4 Enrichment analysis of MACCS fingerprints for CMap-generated lead clusters

Specific bits enriched in MACCS fingerprints of cmap1 and cmap2 leads were selected based on the enrichment scores (ES) defined as the relative frequency of occurrence (p) in the selected set normalized by overall occurrence as follows:

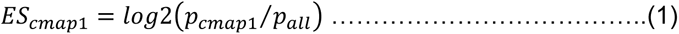

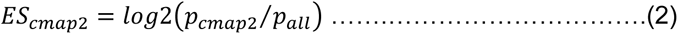

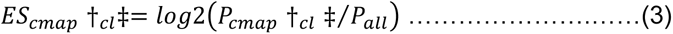

Here, † represents the method of CMap-generated leads i.e. cmap1 or cmap2 and ‡ showed the number of cluster from either of these methods.

### 2.3 Structure-based analysis of CMap-generated leads - Docking experiment

Chemical similarity above helps us to see at enrichments at the ligand-only level. Overall, structure concordance of CMAP leads was assessed using standard molecular docking experiment. To perform docking experiments, a binding site was first identified where the initial docking pose may be created and then the actual docking was performed as follows.

#### 2.3.1 Structure retrieval of primary Target TLR5

An Electron Microscopy (EM) derived model of wild type human TLR5 structure was available in the protein data bank (PDB) (PDBI ID: 3J0A) at the time of starting this work, which had a modest resolution of 26Å (http://www.rcsb.org/structure/3J0A). After, we have already used this structure for the work presented here, AlphaFold tool emerged and provided a complete predicted model. This provided us with a chance to choose the best model of TLR5 between the two. We retrieved the 3D structure of TLR5 from AlphaFold (https://alphafold.ebi.ac.uk/entry/O60602) and compared it with EM model of TLR5. The comparison suggested, a very small variation (RMSD ∼1.27 Å) in the main chain structure of both the models (<3 Å is recommended in **Fusani 2020**) suggesting that the EM and AlphaFold structure of TLR5 are pretty similar to each other. Therefore, we have continued our work with the EM model of TLR5 (see results for details).

#### 2.3.2 Binding site prioritization

We considered two experimentally known leucine-rich repeats (LRR9 and LRR10) region of TLR5 as candidate binding sites from the literature [**Song et al, 2017**]. CastP tool (http://sts.bioe.uic.edu/castp/index.html?2r7g) for cavity detection and sitemap module of Schrödinger Suite for binding site residues prediction in target protein were used. The best site from sitemap prediction was selected according to the sitescore.

#### 2.3.3 Docking experiments

Molecular docking was performed using ligand docking tool in Glide module of the Schrödinger Suite molecular modelling package (version 2019-1). 23-629 amino acid long ectodomain of target protein (TLR5) was prepared using “Protein Preparation Wizard”. In this step, the protein was preprocessed first through bond order and atom type assignment, creation of disulfide bonds, deletion of water beyond the 5 Å and Hetero atoms linked to the target protein then optimized the assigned H-bonds through adjusting the tautomers of ionizable residues (Asn, Gln and His) and finally restrained minimization was performed. Potential binding sites were defined in “Receptor Grid Generation Wizard” to generate the grid around binding site residues in which ligand molecules can dock. CMAP-generated leads were prepared through “LigPrep Wizard” with default parameters. Prepared ligands were docked into potential binding sites with extra precision using Glide to evaluate the best pose of small molecules. All the docking results were evaluated according to the Glide score.

### 2.4 Comparison of TBDD leads with traditional SBDD compounds

One challenge in using docking scores for assessment of favourable binding is the need to have control data sets. To address this question, we used a larger compound library of FDA approved drugs. At the time of data retrieval from DRUGBANK (version 5.1.10) (https://go.drugbank.com/releases/latest#structures/), it consisted ∼2454 FDA approved drugs, A comparison of Glide score between TBDD hits and FDA library compounds was then carried out to determine the favourable gain in binding energy for the selected hits.

### 2.5 Defining high affinity and robust CMap-generated leads

It has been reported that different docking experiments sometimes lead to different prioritized leads and therefore, it would be helpful to performing docking experiments with an additional tool [for example: **Agu et al., 2023**]. For this purpose, we used the popular online tool called **H**igh **A**mbiguity **D**riven protein-protein **DOCK**ing or HADDOCK (version 2.4) which utilizes the information-driven flexible docking approach for the modeling of protein-protein, protein-nucleic acids and protein-ligand complexes [**Zundert et al., 2016**]. In the first round of selection of robust CMap-generated leads, we used another independent tool X-Score (version 1.2.1) which is an empirical scoring function (X-Score = (HPScore + HMScore + HSScore) / 3) that scored the binding affinities for target TLR5 and cmap2 generated leads from both the softwares (Glide and HADDOCK) [**Wang et al., 2002**].

We integrated the outcomes of Glide, HADDOCK and manual analysis of intermolecular interactions, their strengths including hydrogen bonds (H-bonds), hydrophobic interactions (N-bonds) and atom accessibilities by using Ligplot (version 5.1.5) tool [**Wallace et al., 1995**].

### 2.6 Analyzing CMap-generated leads against natural, known and reference ligand of TLR5

As bacterium flagellin *Salmonella enterica* subspecies *enterica serovar* Typhimurium is a 494 amino acid long protein is a well-known natural ligand of TLR5. Its structure has been derived from Electron Microscopy (4Å resolution). We downloaded the 3D coordinates of flagellin from PDB (3A5X) for comparison with CMap-generated leads.

There is only one known non-natural ligand of flagellin namely CBLB502 which is a pharmacologically optimized derivative from *Salmonella enterica* subspecies *enterica serovar* Dublin. Since no structure of CBLB502 is reported or available from AlphaFold database, we modelled the 3D structure of CBLB502 using homology-based approach. To accomplish this task, we first identified the amino acid sequence of CBLB502 (329 amino acid long) in **Burdelya et al. 2008.** Then, this sequence of CBLB502 was used to query the PDB proteins to get the template protein using Protein BLAST tool of NCBI (https://blast.ncbi.nlm.nih.gov/Blast.cgi) then aligning it with target protein sequence using Multiple Sequence Alignment tool Clustal Omega (https://www.ebi.ac.uk/Tools/msa/clustalo/). Finally, on the basis of alignment between target and template, 3D model of CBLB502 was developed and refined it with loop modeling using modeller (version 10.1). Quality of the refined model of CBLB502 was evaluated by ERRAT2 and PROCHECK tools of saves server (https://saves.mbi.ucla.edu/).

Even though, flagellin and CBLB502 have been evaluated well, being large peptide/protein molecules, they do not provide a strong quantitative reference for investigating small molecule binding affinity scores, due to the non-specific differences in their molecular size, higher accessible surface and steric contacts. Manual comparison was therefore used by looking at the interactions of conserved residues ARG90, LEU94, THR102, ASN103, GLU114, GLN117 in the binding pocket.

### 2.7 Experimental validation using ELISA

In order to validate the biological relevance of the prioritized leads, we performed an ELISA assay for them as follows.

#### 2.7.1 Cell Culture

CAL-27 human tongue squamous cell carcinoma cells (RRID:CVCL_1107) were cultured in Dulbecco’s Modified Eagle Medium (DMEM; Sigma, #D5523-10L) supplemented with 10% fetal bovine serum (FBS; Gibco, #10270-106) and 1% antibiotic–antimycotic solution (Sigma, #A5955-100ml). Cells were maintained at 37 °C in a humidified incubator with 5% CO₂. All experiments were performed using cells at passage 8 under identical culture conditions.

#### 2.7.2 ELISA Assay

To determine the production of TLR5 in CAL27 cells treated with selected 9 compounds Fenoterol, Piceatannol, ABT-751, Azacytidine, Cytarabine, Ganciclovir, Kinetin-riboside, Penicillin and Streptozotocin. CAL27 cells (1X106 cells/ml) were seeded in 60mm culture plates and incubated overnight. The cells were treated with different concentrations of compounds for 24h. Cells treated with TLR5 agonist flagellin (Antibodies online, #M1-124021) of concentration 5ng/ml was used as the positive control. Supernatants were collected and centrifuged at 3000rpm for 20min. TLR5 secreted in the culture supernatants were measured using Human Toll-like Receptor 5, TLR5 ELISA kit (BT LAB, #E0374Hu) according to the manufacturer’s protocol. Absorbance was recorded at 450nm using a microplate reader Tecan Spark, and TLR5 concentrations were determined from a standard curve generated by plotting the mean absorbance values against known concentrations of the standard.

### 2.8 Assessment of CMap-generated leads for targeting other proteins in TLR5 pathway

It may be recalled that CMAP protocol does not use structure information of target or ligand and therefore, may select compounds that indirectly activate TLR5. To assess this proposition, we considered downstream proteins of TLR5 signalling pathway as potential other targets of selected hits. We compiled the list of such targets (namely MyD88, IRAK4, IRAK1, TRAF6, TAK1, TAB1, TAB2, MKK6, MKK4, MKK7, IKKA, IKKB, IIKY, P38, JNK, AP1, Ikba, NFKB, IL12, IL6, IL1B and TNFA) and performed a structure based analysis by pairing them with all the selected hots. Each CMap-generated lead was docked to these other-targets using the same procedure as used for TLR5, described above. Glide based docking scores for CMap-generated leads towards TLR5 and other targets were then analyzed using standard statistical tests.

## 3 RESULTS AND DISCUSSION

### 3.1 Generating CMAP leads

#### 3.1.1 DEGs in flagellin activation of TLR5

Scatter plot between the average expression values for control and flagellin activated samples from GSE923 series of GEO (22283 “probes” leading to 13516 genes) is shown in Supplemntary **Figure SF1**. Overall, 49 genes (21 upG or 28 downG) and 21 genes (upG) were used to run corresponding cmap1 and cmap2 queries (See **Supplementary Table ST1 for a detailed list**).

#### 3.1.2 Enriched KEGG pathways associated with DEGs

Although, we cannot directly establish the role of each gene in TLR5 activation, the presence of CCL20, PTX3, CXCL6, CXCL1, CXCL2, IL-8 and CYP1A1 genes was found to be consistent with the analysis presented in the original study of GSE923 reporting the immune responses by TLR5 upon flagellin activation [**Cobb et al., 2004**]. Further plausibility of these DEGs for this phenotype was ascertained using pathways enrichment analysis using SRplot (see Methods and Supplementary **Figure SF2(B**)). Of the top 10 significantly enriched KEGG pathways (FDR < 0.05), several were confirmed to be immune related. These KEGG pathways include Viral protein interaction with cytokine and cytokine receptor (p-value 6.69E-07), Cytokine-cytokine receptor interaction (p-value 3.05E-06), Rheumatoid arthritis (p-value 1.10E-05), IL-17 signaling pathway (p-value 1.16E-05), Chemokine signaling pathway (p-value 2.94E-05), TNF signaling pathway (p-value 2.96E-05), Epithelial cell signaling in Helicobacter pylori infection (p-value 7.29E-05), NF-kappa B signaling pathway (p-value 3.38E-04), Legionellosis (p-value 8.20E-04), and Complement and coagulation cascades (p-value 2.69E-03).

Interestingly, the Viral protein interaction with cytokine and cytokine receptor pathway is highly enriched in comparison to other pathways. It may be noted that flagellin is a potent immunogenic agent and widely explored as a carrier for vaccine development against bacterial, viral, and parasitic diseases [**Mizel and Bates, 2010; Hajam et al., 2017; Zhao et al, 2022**]. Pathways enlisted in databases may therefore be informative of these well-studied associations. Another enriched pathyway namely Complement and coagulation cascades pathway is known to have a significant role in establishing the bidirectional cross-talk between TLR pathways (mainly involves TLR1, TLR2, TLR4, TLR5 and TLR6) and complement system to prevent the excessive inflammatory responses [**Krauss et al., 2000]**. Other enriched pathways are also strongly supportive of the TLR5-mediated flagellin detection and activation/suppression of various protein, transcription factor NF-kappa B, pro-inflammatory cytokines, various kind of interleukins and chemokines involvment in inflammatory disorder, an anti-tumor protection and programmed cell death in the pathogenesis of infectious diseases [**Zeng et al., 2006; Carvalho et al., 2011; Hajam et al., 2017; Losol et al., 2023**].

Thus, the pathway enrichment analysis results support the key assumption in this work that the gene set selected is associated directly or indirectly with TLR5 activation by flagellin and may be confidently used for generating leads with CMAP queries.

#### 3.1.3 CMap-generated leads for flagellin mimics

On the submission of DEGs to CMAP tool, we found a total 80 and 332 CMap-generated leads at 0.9 threshold of CMap score from cmap1 and cmap2 methods respectively (see **Supplementary Table ST1**). We try to validate if these hits, derived solely based on GE signatures are also compatible with structure-based scoring as follows.

### 3.2 Validation of CMAP leads using shared enriched patterns

2One of the core foundations in structure-based drug design is that similar compounds share similar properties and imply similar mode of action against their targets [**Campillos et al., 2008, Gortari et al., 2017**]. However, since CMAP based method mimics only the gene expression environment of flagellin activation and does not use ligand or targets structure or interaction information, it is not clear at the outset if the selected compounds do have a shared TLR5 target or alternatively impact its pathway-level expression in an indirect manner. To gain insights into the mechanistic basis of selections in cmap1 and cmap2 hits, we performed two levels of analysis namely (1) identification of enriched fragments in CMap-generated leads and (2) clustering of leads based on full fragment signature. Results of these analyses are presented below.

#### 3.2.1 Enriched fragments in CMap-generated leads

Overall relative frequency of each fragment in cmap1 and cmap2 leads was compared with the background relative frequencies of CMAP database library (all compounds in CMAP against which leads were searched). **Table 1** lists out the selected fragments enriched in these hits. We observed that some of the MACCS fingerprints were enriched in selected lead compounds, although no single compound was found to be decisively representative of flagellin response. For example **Table 1 [A]** Bit26, Bit13, Bit24 and Bit23 were over-represented and Bit52 was depleted in cmap1 leads. Three of the top 5 enriched fragments have a nitrogen atom in them? In addition, MACCS fragment called #CHARGE was highly enriched indicating that flagellin mimics may have an electrostatic interaction with their targets.

**Table 1:**
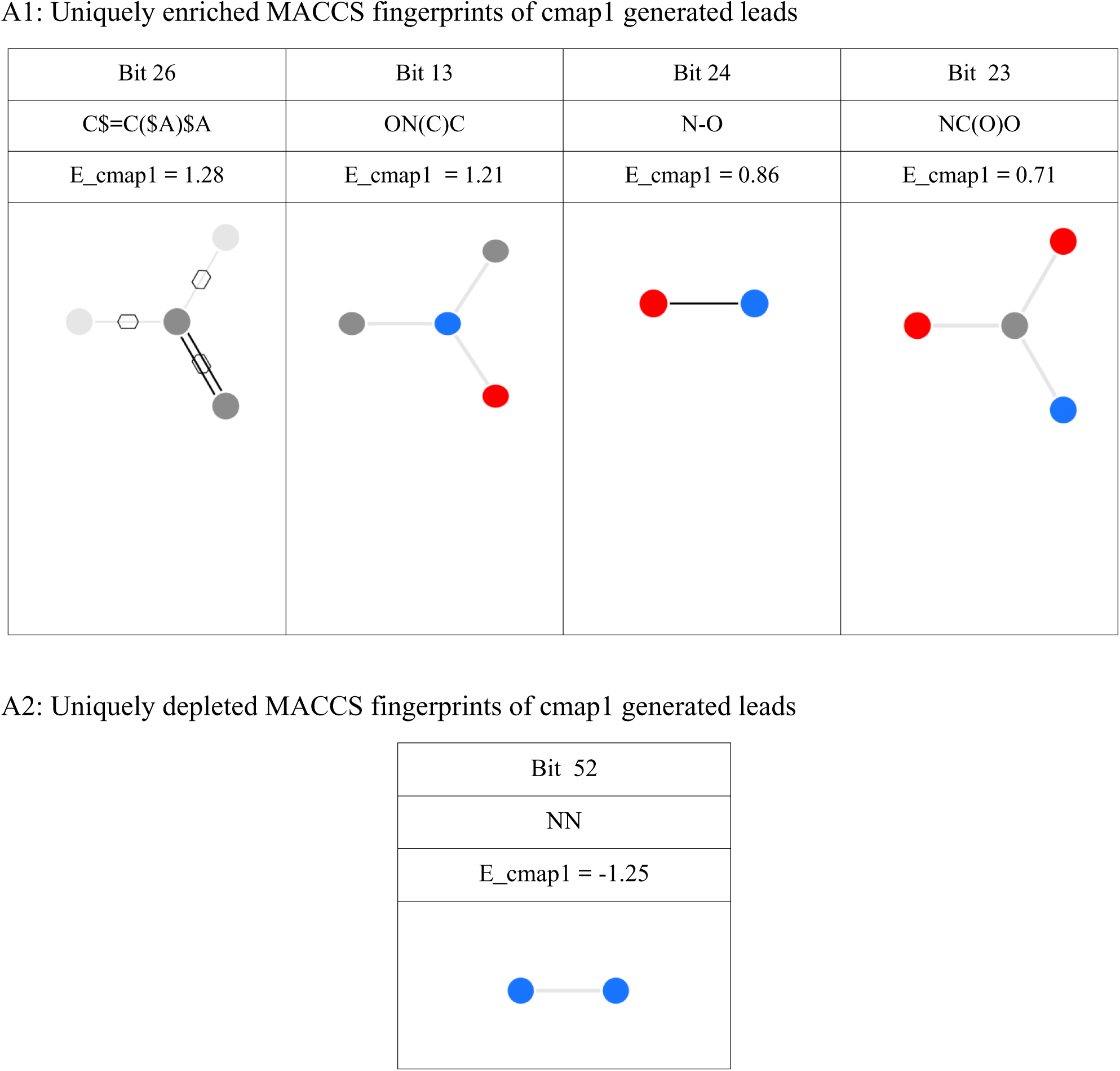
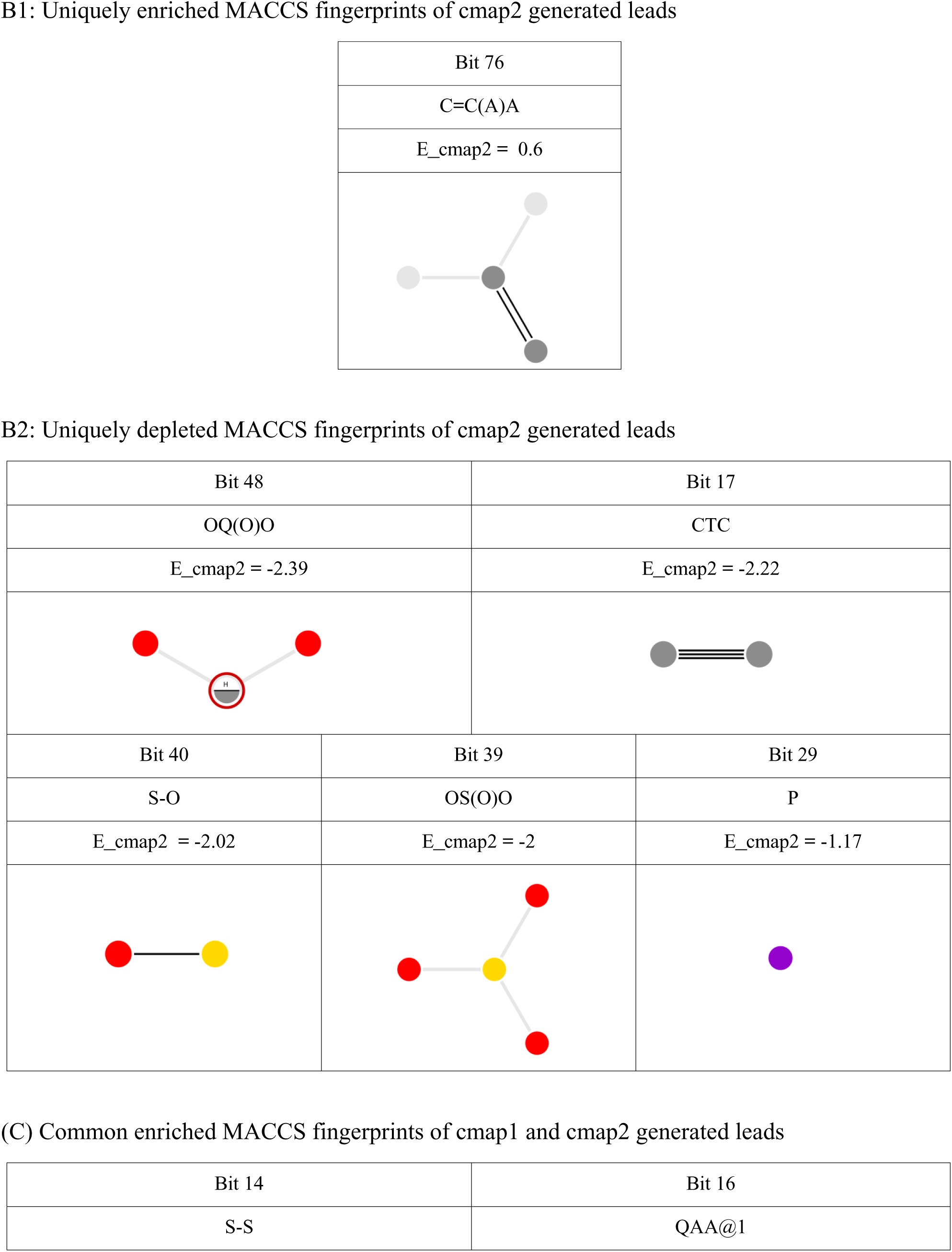
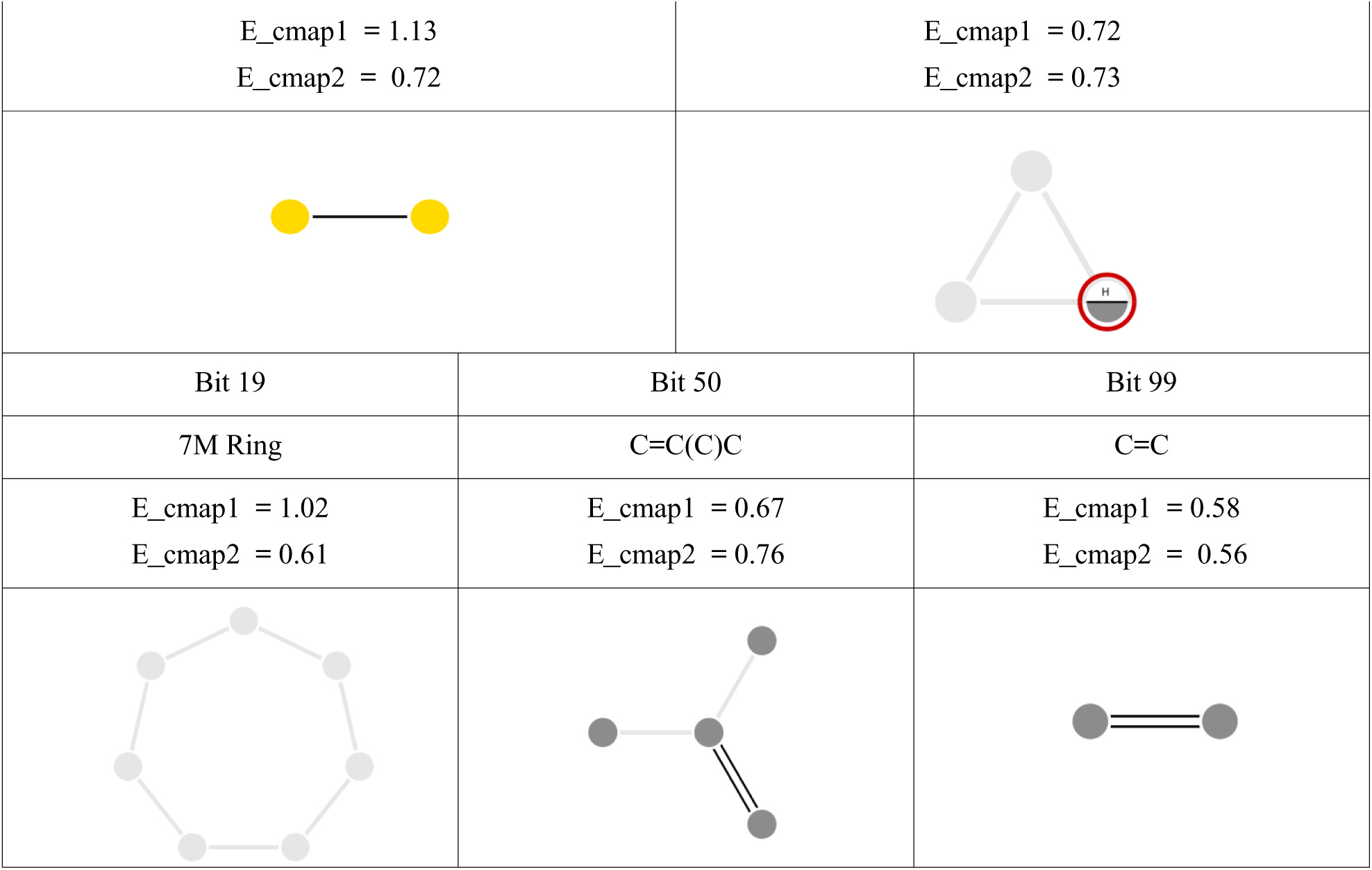
List of enriched and depleted fragments in cmap1 and cmap2 hits for flagellin activated expression profiles. (A) and (B) for uniquely generated cmap1 and cmap2 leads and (C) for common between cmap1 and cmap2 generated leads. *Note that enrichment scores are on log scale and a score of 1.0 means 2-fold increase. Negative log values represent depletion of corresponding fragments in the CMAP hits. The enriched MACCS fingerprints were plotted using the SMARTS.plus online tool. Light grey sphere and line represent arbitrary atom connected by any type of bond. Red circles with hydrogen denote any non-C or non-H atom (only hetero atoms) that should belong to any valid periodic table element. Hexagons along with bonds denote that this bond is within a cyclic system. Two dark grey lines represented double bond between carbon-carbon atom. Dark grey, red, blue, yellow and purple sphere indicate carbon, oxygen, nitrogen, sulpher and phosphorus atom respectively*.

**A1: Uniquely enriched MACCS fingerprints of cmap1 generated leads**
| Bit 26 | Bit 13 | Bit 24 | Bit 23 |
| --- | --- | --- | --- |
| C\$=C(\$A)\$A | ON(C)C | N-O | NC(O)O |
| E_cmap1 = 1.28 | E_cmap1 = 1.21 | E_cmap1 = 0.86 | E_cmap1 = 0.71 |

**A2: Uniquely depleted MACCS fingerprints of cmap1 generated leads**

| Bit 52 |
| --- |
| NN |
| E_cmap1 = -1.25 |

B1: Uniquely enriched MACCS fingerprints of cmap2 generated leads

|  |  |  |  |
| --- | --- | --- | --- |
| Bit 48 |  | Bit 17 |  |
| OQ(O)O |  | CTC |  |
| E_cmap2 = -2.39 |  | E_cmap2 = -2.22 |  |
| Bit 40 | Bit 39 |  | Bit 29 |
| S-O | OS(O)O |  | P |
| E_cmap2 = -2.02 | E_cmap2 = -2 |  | E_cmap2 = -1.17 |

(C) Common enriched MACCS fingerprints of cmap1 and cmap2 generated leads
|  |  |
| --- | --- |
| Bit 14 | Bit 16 |
| S-S | QAA@1 |

|  |  |  |
| --- | --- | --- |
| $E_{\text{cmap1}} = 1.13$<br>$E_{\text{cmap2}} = 0.72$ | | $E_{\text{cmap1}} = 0.72$<br>$E_{\text{cmap2}} = 0.73$ |
| Bit 19 | Bit 50 | Bit 99 |
| 7M Ring | C=C(C)C | C=C |
| $E_{\text{cmap1}} = 1.02$<br>$E_{\text{cmap2}} = 0.61$ | $E_{\text{cmap1}} = 0.67$<br>$E_{\text{cmap2}} = 0.76$ | $E_{\text{cmap1}} = 0.58$<br>$E_{\text{cmap2}} = 0.56$ |

Interestingly the enriched fragments for cmap2 leads are quite different from cmap1 hits. **Table 1 [B]** shows that Bit48 , Bit40 , Bit39 and Bit 29 which was enriched in cmap2 hits, while many were actually depleted. The depleted fragments contained oxygen, sulfur and phosphorus atoms in contrast to nitrogen related fragments found enriched in cmap1 leads. Bit14 fragment SS is the with sulfur-based fragments and Bit50 and Bit99 with a C=C fragments and the 7-member ring based fragment Bit19 were enriched in both cmap1 and cmap2 (see **Table 1 [C])**.

Thus, the enrichment analysis of the fragments provides not only the confidence that the cmap1 and cmap2 hits are not structurally random but also give useful insights into potential roles of their atomic contents. We also note that the enrichment scores are not really decisive individually, suggesting that several atomic subgroups are involved in flagellin-like activation of TLR5 either in a cooperative or alternative manner.

Since, single atomic fragments were only weakly enriched; a better signal could be detected by clustering overall fragment signatures of selected hits. Thus, we next explore if the selected leads come from distinct clusters of atomic signatures.

#### 3.2.2 Clustering of lead compounds based on fragments based fingerprints

Using a simple clustering based approach to identify groups of lead compounds with similar chemical signatures in cmap1 and cmap2 generated leads, we found that cmap1-generated leads formed two cluster of 36 and 44 leads while cmap2 generated leads were grouped into three clusters of 96, 41 and 181 leads respectively (see **Supplementary Table ST2**). Enrichment analysis of the fragments in each clusters is presented in **Supplementary Table ST3**. We observe that the fragment-wise enrichment in clusters is better than overall cmap1 and cmap2 hits analyzed independently, suggesting that identified leads come from specific groups of ligand molecules, perhaps providing alternative modes of binding. For example for cmap1 hits, cluster1 enrichment scores are as high as 2.36 and 2.21 for Bin 13 and Bin 26 while in cluster 2 has enrichment of 2.92 and 1.99 respectively for Bit 16 and Bit 50. None of the fragments was enriched by this degree in the overall sets of fingerprints of cmap1 and cmap2 leads, discussed in the previous section.

To gain further insights into mechanistic basis of identified hits, we investigate molecular interactions of hits with their target using molecular docking experiments.

### 3.3 Validation of CMap-generated leads using direct interaction analysis

#### 3.3.1 Modeling target TLR5 structure and binding sites

Analysis of chemical fragments and their clusters in cmap1 and cmap2 leads, discussed above suggests some interesting patterns in their fragment-based signatures. A better and more direct structure based validation of these compounds can be carried out by a direct molecular docking experiment, which requires a three-dimensional structure data and identification of potential binding pockets . As mentioned in Methods, we have used CryoEM-based structure of TLR5 for our modeling experiments. Resulting structure of eco-domain of TLR5, which is the main target for flagellin is shown in Supplementary **Figure SF2**, together with its prioritized binding region as discussed in Methods. Consensus residues for this potential binding site are residue ID: 233, 235, 236, 237, 240, 243, 246, 247, 250, 258, 260, 261, 262, 263, 264, 266, 267, 268, 270, 271, 272, 273, 281, 282, 285, 293, 311. With this information, we can now proceed to investigate interaction energies of various leads using molecular docking.

#### 3.3.2 Molecular docking experiments

Molecular docking was conducted on generated leads and also for an entire list of 2454 FDA-approved drugs as the control set. A detailed comparison of docking score and Cmap score for cmap1 and cmap2 leads are shown in **Supplementary Table ST4.** In summary, we observed that 56-62% of the compounds successfully docked on TLR5. The FDA control, cmap1 and cmap2 compounds did not differ much as far as the overall proportion of successfully docked compound is concerned (p-value using a chi-squared test was 0.56 and 0.12 respectively when compared by expected counts take from control and observed counts from cmap1 and cmap2). The specificity emerged in the top scoring complexes (see **Table 2) i.e.** when we look at the number of compounds that bind TLR5 tightly (glide score < -6.0 kcal/mol), we find that 18 out of 46 cmap1 and 63 out of 185 cmap2 compounds showed tight binding compared to 336 out of 1531 in the case of FDA control. Thus, in this comparison cmap1 and cmap2 were found to be enriched in tightly binding compounds with a p-value of 0.0129 and 0.0004 respectively. This suggests strong evidence in favour of the selectivity of cmap1 and cmap2 in reference to a randomly selected compound from FDA approved drugs. This is particularly impressive considering that no structural or direct binding specificity information was used in selecting these leads.

**Table 2.** Summary of docking experiments performed on TLR5 structure using cmap1 and cmap2 leads and FDA compounds as a control. *P-values have been computed by using FDA compounds to compute expected counts in each category and applying a Chi squared test. While the docking success rate in the FDA, cmap1 and cmap2 compounds have no significant statistical difference, high scoring compounds are statistically significantly enriched in both cmap1 and cmap2 selected leads*.

| **Compound list** | **No of compounds** | **No of uniquely docked compounds with TLR5 in Glide** | **Success rate in docking** | **Compounds with glide score above -6 kcal/mol** | **p-value (docking success)** | **p-value (high scoring compounds)** |
| --- | --- | --- | --- | --- | --- | --- |
| FDA control | 2454 | 1531 | 62% | 336 |  |  |
| cmap1 | 80 | 46 | 58% | 18 | 0.5799 | 0.0129 |
| cmap2 | 332 | 185 | 56% | 63 | 0.1242 | 0.0004 |

### 3.4 Prioritization of CMap-generated leads

Having established that CMAP leads are enriched in fragment-based signatures and provide high specificity against other FDA approved drugs, we now move on to further integrate the information from molecular interaction analysis and CMAP scores to prioritize leads for experimental validation. For this purpose, we analyzed all the tightly docked cmap2 generated leads first on the basis of their binding free energy through X-Score and the manually examined the plausible inter-molecular interactions including hydrogen bonds (H-bonds) and hydrophobic interactions (N-bonds) by Ligplot tool as follows.

#### 3.4.1 Interaction analysis of TLR5 complex with cmap hits

Detailed outcomes of docking and interaction analysis are provided in (See **Supplementary Table ST5**). All the tightly docked cmap2 generated leads were predicted to be strong binders to target TLR5 since their scores of binding affinities from X-Score were also < -6.0 kcal/mol. Beside this, 13 out of 63 cmap2-generated leads from Glide Ligplot and 31 out of 63 cmap2 genertaed leads from HADDOCK Ligplot showed strong interactions with TLR5 through H-bonds (considering minimum 3 H-bonds interactions between TLR5 and cmap2 generated leads were strong interaction). Non-bonded (N-bonds) interactions which involve Van der Waals and electrostatic interactions were also considered in selection of Cmap-generated leads. We prioritized 9 tightly docked cmap2-generated leads **(see Table 3)**, selected by a consensus of both the interactions (Ligplot of Glide and HADDOCK both). Looking back at the clusters of compounds based on molecular fragments discussed above, 9 robust CMap-generated leads provided sufficient coverage of chemical diversity (2 from cluster1, 1 each from cluster2 and 6 from cluster 3). Graphical representation of intermolecular interactions of robust CMap-generated leads and TLR5 are shown in **Figure 2**.

**Figure 2:**
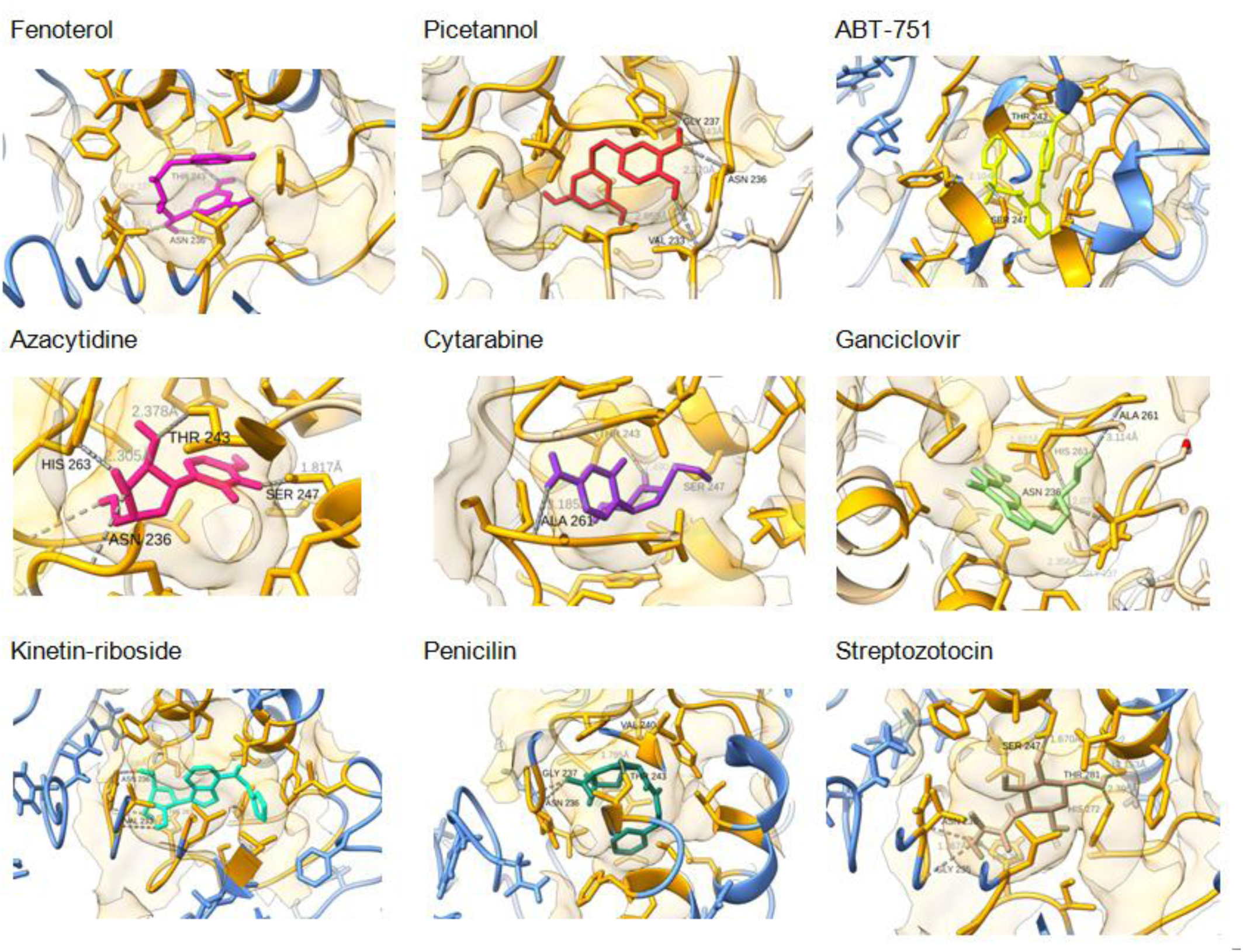
Intermolecular interactions of representative CMap-generated leads with TLR5. This figure shows all the H-bond interactions between robust CMap-generated leads and TLR5 which fall below 3 Å. The minimum 3 H-bond interactions between CMap-generated leads and TLR5 were showed by both Glide module and HADDOCK tool in each complex.

**Table 3.** Robust CMap-generated leads and its structural validation. Selecting robust CMap-generated leads using TBDD. A comparison based on binding free energy, X-Scores and intermolecular interactions (H-bonds and N-bonds) by Ligplot between robust CMap-generated leads and reference molecules with target TLR5 after docking experiment using Glide module and HADDOCK tool. Here ChemScene id is abbreviated by CS

| Robust CMap-generated leads | CMap scores | cmap2 cluster no. | Glide Score (kcal/mol) | Glide Xscore (kcal/mol) | Glide Ligplot H-bond counts | Glide Ligplot H-bonds | Glide LigPlot N-bonds counts |
| --- | --- | --- | --- | --- | --- | --- | --- |
| Reference molecule | None | None | -6.99 | -6.95 | 3 | T243, N236, V233 | 41 |
| Fenoterol (CS-5285) | 99.98 | c1 | -8.31 | -8.55 | 3 | H272, T243, N236 | 67 |
| Piceatannol (CS-5657) | 96.97 | c1 | -7.24 | -8.51 | 5 | G237, 2N236, G235, V233 | 46 |
| ABT-751 (CS-0495) | 99.86 | c2 | -6.99 | -9.11 | 5 | N273, H272, H263, S247, T243 | 82 |
| Azacytidine (CS-1287) | 99.52 | c3 | -6.91 | -6.96 | 3 | H263, T243, N236 | 34 |
| Cytarabine (CS-2177) | 96.67 | c3 | -6.67 | -7.06 | 4 | A261, 2S247, T243 | 23 |
| Ganciclovir (CS-2014) | 99.57 | c3 | -7.76 | -7 | 5 | H263, A261, G237, N236, V233 | 34 |
| Kinetin-riboside (CS-6377) | 99.95 | c3 | -9.01 | -8 | 3 | 2N236, V233 | 77 |
| Penicillin (CS-3873) | 92.46 | c3 | -6.30 | -8.89 | 3 | T243, G237, N236 | 78 |
| Streptozotocin<br>(CS-1828) | 99.84 | c3 | -8.14 | -7.16 | 4 | H272, H263, S247, N236 | 47 |

B: Comparison based on HADDOCK tool
| **Robust CMap-generated leads** | **CMap scores** | **cmap2 cluster no.** | **Haddock Score (kcal/mol)** | **Haddock Xscore (kcal/mol)** | **Haddock Ligplot H-bond counts** | **Haddock Ligplot H-bonds** | **HADDOCK LigPlot N-bonds counts** |
| --- | --- | --- | --- | --- | --- | --- | --- |
| Reference molecule | None | None | None | None | 6 | R306, A261, S247, F238, V233, L208 | 78 |
| Fenoterol<br>(CS-5285) | 99.98 | c1 | -36.4 | -7.49 | 4 | H262, A261, G237, V233 | 71 |
| Piceatannol<br>(CS-5657) | 96.97 | c1 | -31.2 | -8.07 | 3 | H263, G237, N236 | 67 |
| ABT-751<br>(CS-0495) | 99.86 | c2 | -28.7 | -7.68 | 4 | G237, N236, G235, V233 | 73 |
| Azacytidine<br>(CS-1287) | 99.52 | c3 | -29 | -7.08 | 3 | S247, T243, V233 | 43 |
| Cytarabine<br>(CS-2177) | 96.67 | c3 | -27.7 | -7.1 | 6 | N273, 3G237, N236, V233 | 50 |
| Ganciclovir<br>(CS-2014) | 99.57 | c3 | -32.8 | -7.15 | 6 | N273, H263, A261, T243, N236, G235 | 51 |
| Kinetin-riboside<br>(CS-6377) | 99.95 | c3 | -40.9 | -8.07 | 5 | 2H263, 2G237, G235 | 79 |
| Penicillin<br>(CS-3873) | 92.46 | c3 | -37.4 | -7.29 | 4 | T243, V240, G237, N236 | 56 |
| Streptozotocin<br>(CS-1828) | 99.84 | c3 | -30.6 | -7.08 | 8 | H262, A261, I259, N236, 2G235, S234, V233 | 72 |

#### 3.4.2 Comparison with natural ligand, known ligand and reference molecules

Having identified 9 top candidates for experimental validation, we also compared them with their known ligands of TLR5. Other than the natural ligand flagellin and the only other known ligand for TLR5 is CBLB502 and both of them are much larger than our leads, making a direct comparison of binding energies challenging. We nonetheless put similar docking experiments on these compounds in perspective (see **Table 4**). The 3D structure of CBLB502 which was modelled by homology modeling tool Modeller (version 10.1) for this purpose. We found that the 84.6 % sequence of CBLB502 (protein length = 329 amino acid) was identical to Salmonella enterica subspecies enterica serovar Typhimurium R-type straight flagellin (protein length = 506 amino acid and PDB id = 6JY0, Chain A) with Expected value of 2.00E-94. Total 25 models were generated in which best model was identified using DOPE (Discrete Optimized Protein Energy) score. Further, the lowest DOPE score model was evaluated by ERRAT2 tool of saves server, highlighting the three loop regions (at position 39-47, 138-144 and 158-164) were exceeding the error value of rejecting regions (above 99%) (see **Supplementary Figure SF3**) and showed the overall quality factor of model is 84.3. After refining the above mentioned loops, the best model out of 10 loops refined models was selected on the basis of DOPE score and evaluated again using ERRAT2 and PROCHECK tools. We observed that the overall quality factor of the best loop refined model of CBLB502 was improved from previous model which is 92.5 (**see Supplementary Figure SF3(B)**). In addition to this, PROCHECK evaluation suggested that the 93.4% of residues of CBLB502 model were found in the most favored regions, while none were found in disallowed regions (see **Supplementary Figure SF3(C)**). Final model of CBLB502 was visualized through chimeraX and compared with flagellin which showed a minor deviation of 0.913 Å in their backbone structure (see **Supplementary Figure SF3(D)**). All the above mentioned results of homology modeling suggested that the our 3D model of CBLB502 is good enough to dock to target TLR5 and can predict their binding free energy and inter-molecular interactions with target TLR5.

**Table 4:**
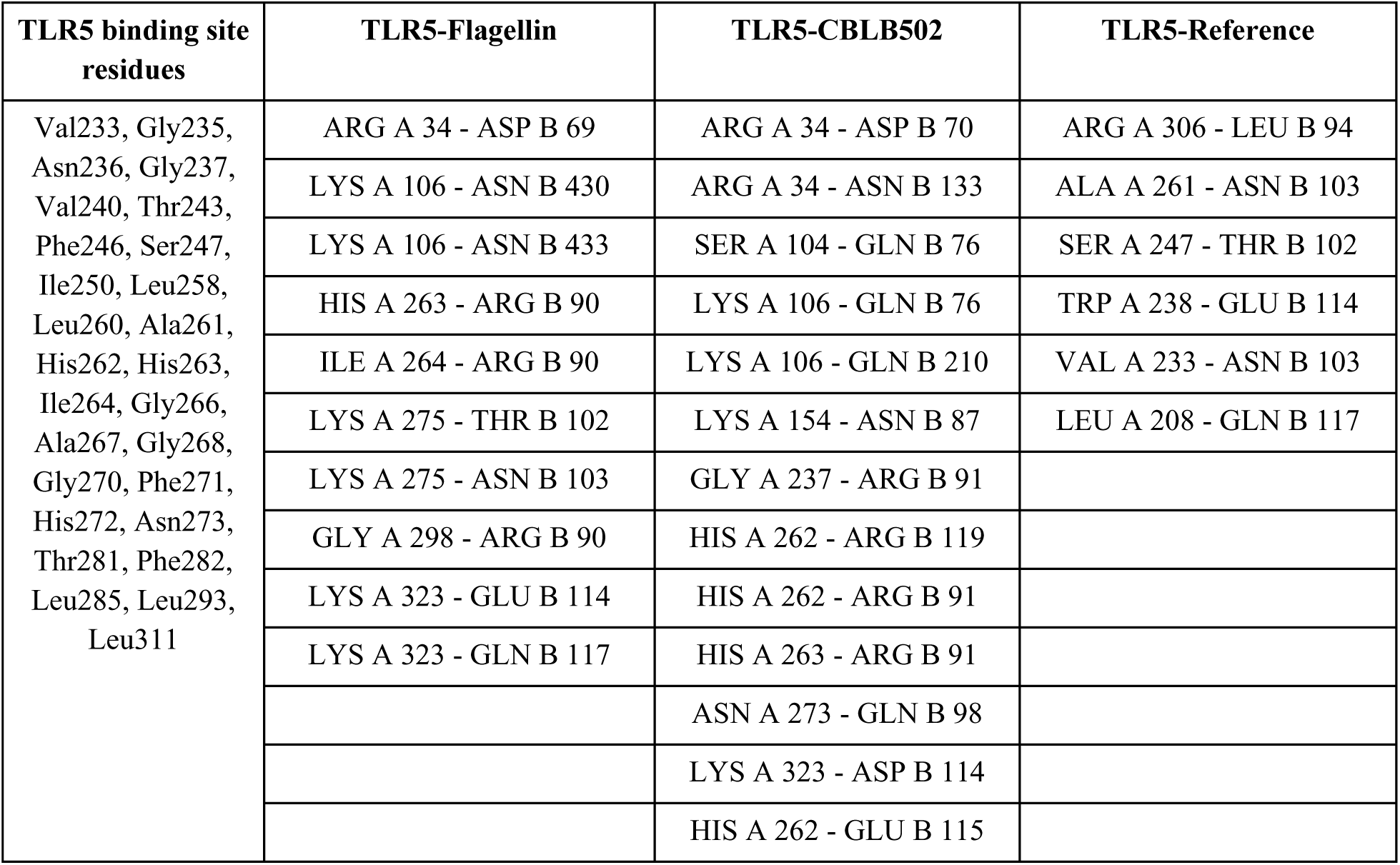
Comparison of CMap-generated leads with natural, known and reference ligand of TLR5. Comparison of inter-molecular interactions between natural ligand: flagellin, known ligand: CBLB502 and reference molecule with the binding pocket residues of TLR5 after docking using HADDOCK

After modeling a good quality structure of CBLB502 and the available structure of flagellin (PDB id: 3A5X), we docked both the structures of CBLB502 and the flagellin to the target TLR5 separately using the HADDOCK server. Then, we looked into the H-bond interactions in both complexes (TLR5-flagellin and TLR5-CBLB502) using the chimeraX tool (**Supplementary Figure SF4).**

Furthermore, X-Score’s binding free energy appears to be a suitable parameter for this comparison as Glide docked complex of TLR5 and reference molecule consisting of (ARG90, LEU94, THR102, ASN103, GLU114, GLN117 from flagelin) give an X-score of -6.95 kcal/mol. In comparison with reference molecule, all the robust CMap-generated leads also showed comparable but higher X-Scores upon after docking with Glide as well as HADDOCK, suggesting these leads may be potent binders of TLR5 with the same strength as the latter’s known ligands (**Table 3**).

### 3.5 Experimental assessment of TLR5 differential regulation by prioritized leads

To assess the modulatory effects of selected small molecules on TLR5 expression, cells were treated with 9 leads listed in **Table 3** namely Fenoterol, Piceatannol, ABT-751, Azacytidine, Cytarabine, Ganciclovir, Kinetin-riboside, Penicillin-G and Streptozotocin. TLR5 expression levels were subsequently quantified and compared to untreated control samples. **Figure 3** shows the outcomes of these results where TLR5 activation is confirmed in all the 9 leads but the concentration dependence is somewhat different. For example, treatment with five of these 9 leads namely Cytarabine, Azacytidine, Penicillin-G, Streptozotocin and Piceatannol led to notable increase in TLR5 expression levels relative to the control group with their increasing concentrations (See **Figure 3A**). Cytarabine treatment led to a concentration-dependent increase in TLR5 expression, which may reflect a cellular stress response or compensatory immune priming due to chemotherapy-induced cytotoxicity **[Ersvaer et al., 2015]**. Similarly, treatment with Azacytidine resulted in a dose-dependent upregulation of TLR5 expression. This can potentially be attributed to its hypomethylating action, causing reactivation of silenced TLR5 gene expression via promoter demethylation **[Tsuji-Takayama et al., 2004]**. While Penicillin-G is not a known direct agonist of TLR5, the observed increase may reflect an indirect modulation of immune receptor expression via low-level of inflammatory signaling **[Moore et al., 2003]**. Penicillin-G increases IL-6 **[Yousef et al., 2021]** and decreases IL-10 **[Volk et al., 2020]** in some models due to cytokine modulation which further may stimulate TLR5 via NF-κB. Streptozotocin, a nitrosourea compound, treatment induced TLR5 expression, which is likely mediated by oxidative stress and NF-κB activation [**Bathina et al., 2017**]. These findings highlight possible intersection between cytotoxic stress and innate immune receptor regulation. Piceatannol, a known polyphenolic compound with anti-inflammatory properties, may enhance TLR5 expression via redox-sensitive transcriptional pathways [**Lee et al., 2010**].

**Figure 3:**
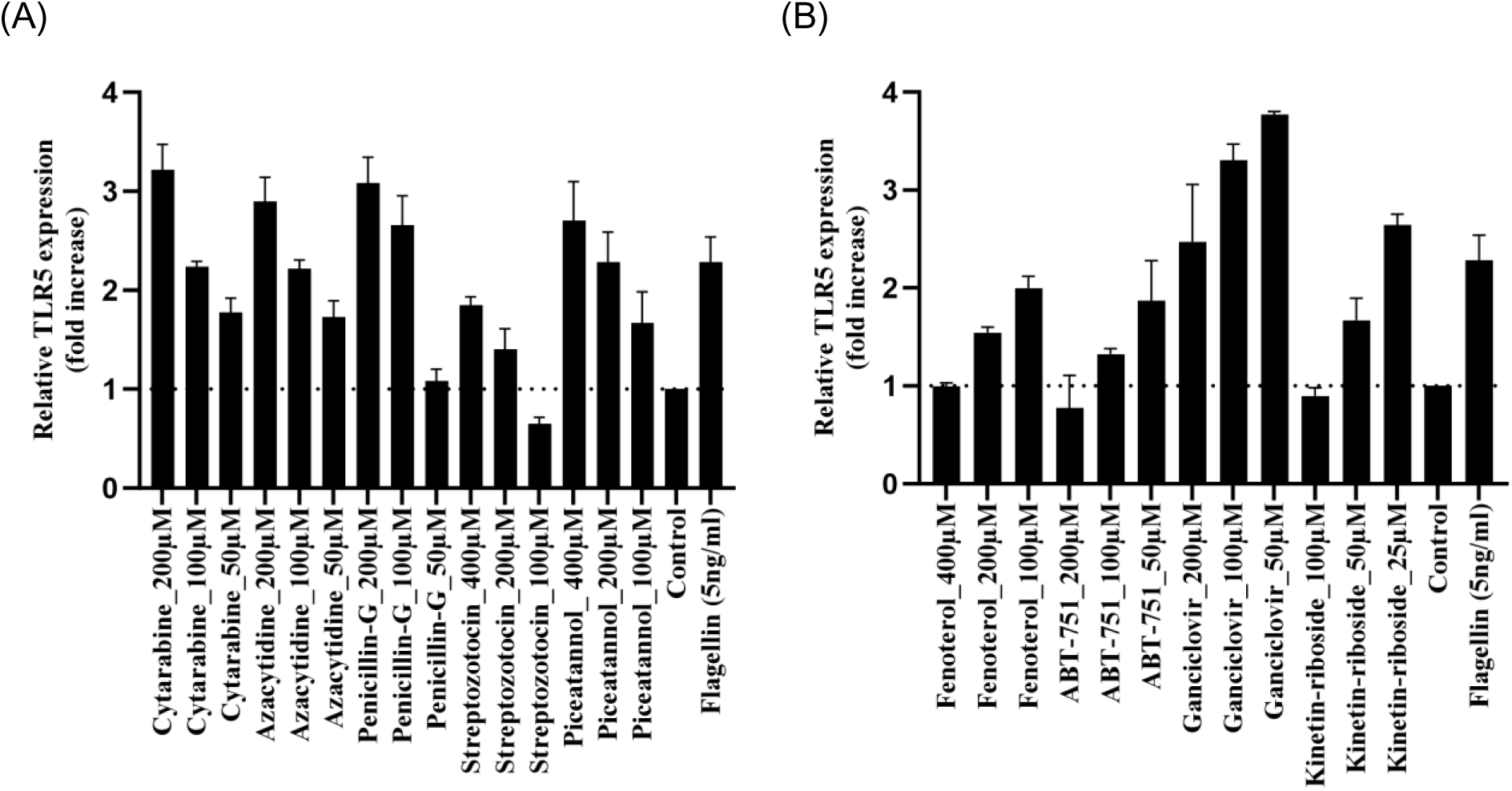
Detection of TLR5 expression in CAL27 cells in response to CMap-generated leads. (A) CAL 27 cells were incubated with the CMap-generated leads: Cytarabine (200µM, 100µM, 50µM), Azacytidine (200µM, 100µM, 50µM), Penicillin (200µM, 100µM, 50µM), Streptozotocin (400µM, 200µM, 100µM), Piceatannol (400µM, 200µM, 100µM) and flagellin (5ng/ml) for 24h. (B) CAL 27 cells were incubated with the CMap-generated leads: Fenoterol (400µM, 200µM, 100µM), ABT-751 (200µM, 100µM, 50µM), Ganciclovir (200µM, 100µM, 50µM), Kinetin-riboside (100µM, 50µM, 25µM), and flagellin (5ng/ml) for 24h. Bar graph representing the relative expression levels of TLR5 with respect to the untreated control. Fold change is calculated by the signal of the treated sample divided by the signal of the control. If the fold change is >1, TLR5 expression is upregulated whereas fold change <1 is taken as downregulated. All values are expressed as mean ± SEM.

Four of the prioritize leads (**Figure 3B**) show a somewhat different concentration dependence of TLR5 activation. Of these, Fenoterol is a selective β2-adrenergic receptor (β2-AR) agonist commonly used as a bronchodilator in asthma and COPD [**Svedmyr 1985**]. Beyond its respiratory effects, fenoterol modulates immune responses via β2-adrenergic signaling, which is known to have anti-inflammatory and immunosuppressive properties. Since Fenoterol treatment resulted in a dose-dependent *decrease* in TLR5 expression, it is consistent with β2-adrenergic receptor–mediated immunosuppression [**Alonso and Iñigo-Marco, 2019**]. This downregulation may be attributed to cAMP/PKA signaling, which suppresses NF-κB activation, a key regulator of TLR5 transcription [**Wang et al., 2016**]. These findings highlight the potential of Fenoterol to modulate innate immune recognition pathways in addition to its bronchodilatory effects. Similarly, treatment with ABT-751 led to a dose-dependent *reduction* in TLR5 expression. This decline is likely attributable to ABT-751–mediated disruption of microtubule dynamics [**Wei et al., 2016**], which impairs TLR5 trafficking and downregulates its gene expression. Additionally, ABT-751–induced mitotic arrest and cytotoxic stress may contribute to immune desensitization by dampening innate immune receptor expression [**Dehghanian et al., 2021**]. A concentration-dependent *decrease* in TLR5 expression was also observed upon treatment with Ganciclovir. This effect is likely mediated by its anti-proliferative activity and associated immune-suppressive mechanisms [**Battiwalla et al., 2007**]. Finally, the treatment with increasing concentrations of Kinetin Riboside led to a marked, dose-dependent *decrease* in TLR5 expression likely attributed to KR-induced mitochondrial stress, apoptotic signaling [**Choi et al., 2008**] and suppression of NF-κB–mediated transcription [**Tiedemannet al., 2008**]. These findings collectively underscore the unexpected immunomodulatory potential of several pharmacological agents and suggest possible off-target or secondary roles in modulating innate immune sensors like TLR5.

As a first step towards understanding the possibility of selected hits also interacting with other members of TLR5 pathway, we repeated our docking experiments on them as follows.

### 3.6 Potential interactions of leads with other pathway proteins

We identified 24 targets closely associated to TLR5 pathway listed in **Supplementary Table ST6**. The atomic coordinates and their selected binding pockets are provided in **Supplementary Table 3.** All the cmap1 and cmap2 compounds were docked to all these 24 off-target proteins in a way similar to TLR5 and the docking scores of each of these compounds are shown in **Supplementary Figure SF5 (A).** A statistical comparison between docking scores against TLR5 and members of its signaling pathway shows is shown in **Supplementary Figure SF5 (B)**. Results suggest that these additional pathway proteins have a correlation ranging from the highest value of 0.380 for TAK1 to the lowest 0.136 for MKK6. Interestingly there are no negative correlations, suggesting that the cmap1 and cmap2 compounds selected for TLR5 may also have some hidden similarity with TLR5 and possibly show some weak binding to other pathway proteins. Some of these may provide systems level explanation to experimental outcomes. However, these insights are indicative and a detailed mechanistic basis of observed dose-dependent associations of leads uncovered in this work, will require much more work in the future.

## 4. CONCLUSION

To date, molecular docking or ligand chemical signature are the most widely used methods for novel lead identification. We explored in this work a novel method by combining transcriptome based lead generation with conventional molecular docking and fragment signature enrichment leading upto nine potential leads for further experimentation. All the nine leads were found to successfully mimic activation of TLR5 by its natural ligand. This shows that the proposed framework of selecting leads for experimental validation works efficiently and better mimics the cellular response, inherent in the protocol. However, dose dependence of TLR5 activation in some compounds calls for a close inspection of molecular responses during the experimental validation stage.

## 5. SUPPLEMENTARY MATERIALS

All supplementary data is provided in supplementary-data.pdf

## 6. DECLARATION OF COMPETING INTEREST

The authors declare that they have no known competing financial interests or personal relationships that could have appeared to influence the work reported in this paper.

## 7. AUTHOR CONTRIBUTIONS

A.J.: Full computational implementation and MS draft preparation N.S.: critical analysis and suggestions HH & V.T.: Wet lab experimental validation S.A.: conceptualization, supervision, MS editing and overall experimental design. All authors have read and agreed to the published version of the manuscript.

## 8. FUNDING

This research was supported by a research grant ICMR grant AI-Adhoc/12/2022-AI Cell (awarded to SA; post doctoral fellowship to AJ), DBT grant (no. BT/PR24208/BID/7/801/2017), Bioinformatics Center (DBT-BIC) and National Networking Project (NNPs with IIITD and CSIR-NEIST) from the Department of Biotechnology, Government of India (awarded to SA) and an ICMRGrant no. ICMR-SRF:2020-8554 (Senior Research Fellowship to AJ).

## Notes

### Competing Interest Statement

The authors have declared no competing interest.

