## supplementary for "Transcriptome-based lead generation, ligand- and structure-based prioritization and experimental validation of TLR5-activating molecules"

**Supplementary Figure SF1: Differentially expressed genes (DEGs) for query to CMAP and their pathways enrichment.**

*(A) Scatter plot of DEGs for control vs flagellin active profiles at log2FC thresholds of 2 (B) Enriched pathways upon shortlisted DEGs.*

(A)

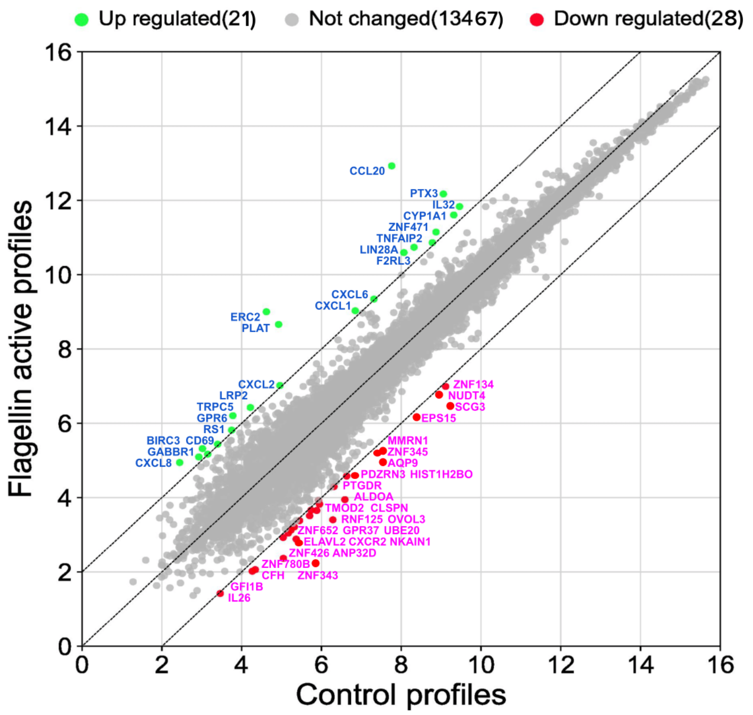


(B)

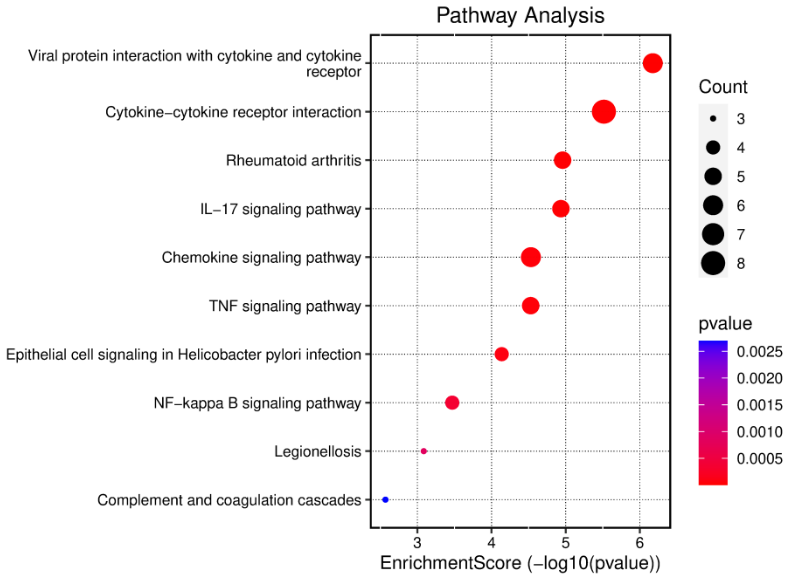

### Supplementary Figure SF2: The 3D structure of the TLR5 (chain A only) inferred from the EM model (3J0A: blue in color).

*(A) A full length structure (extracellular, transmembrane and cytoplasmic domain) of TLR5 (B) Extracellular structure and the inset view of binding pocket of TLR5.*

*Consensus binding pocket was highlighted with an golden color on the surface of TLR5 while its residues were zoomed in and showing the right side of the figure. Two disulfide bonds were represented by red and yellow color respectively. The FUC (l-fucose) and NAG (N-acetylglucosamine) molecules were represented by magenta and green color respectively.*

(A)

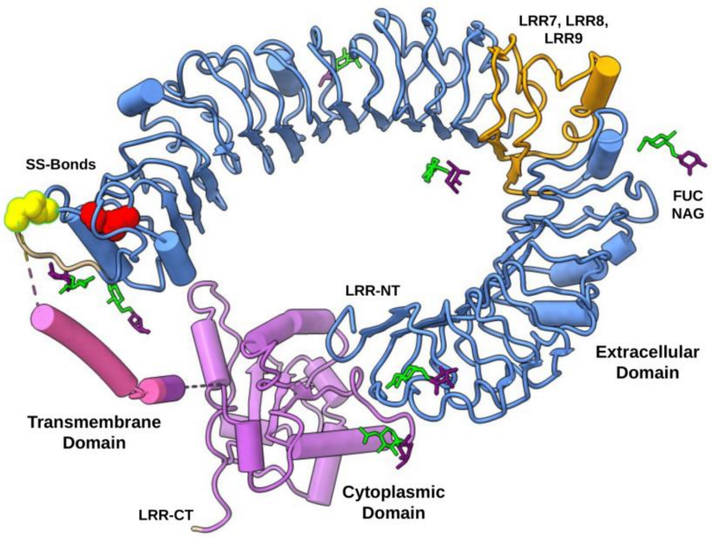


(B)

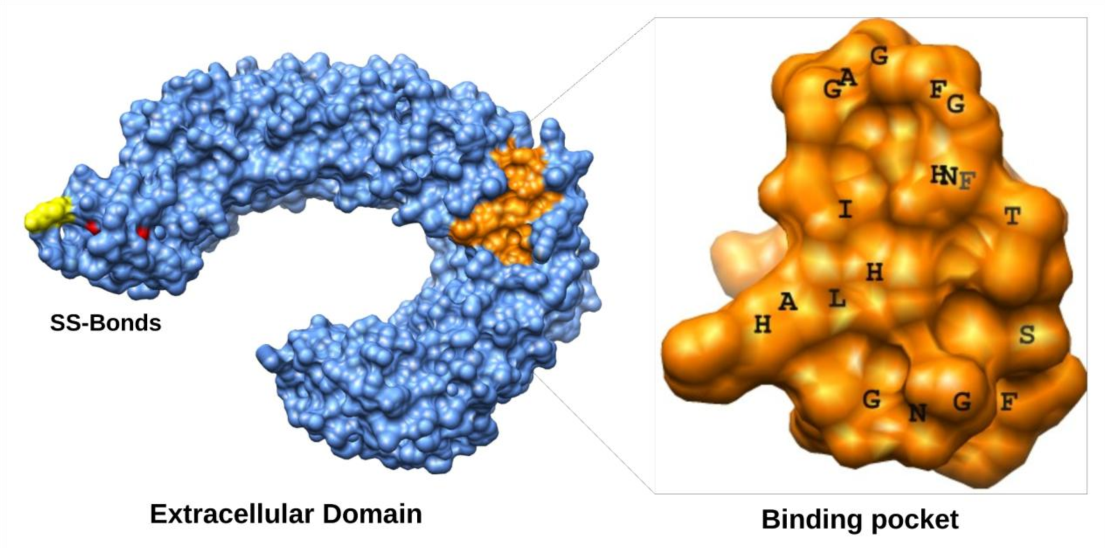

***Supplementary Figure SF3 Structural evaluation of the homology model of CBLB502.***

*This figure explains the overall consistency of the homology-modeled structure of CBLB502. Subplot 12a and 12b highlight the quality score for the CBLB502 model before and after loop refinement using ERRAT2 tools and 12c describes the overall information of the backbone residues using PROCHECK tool of the SAVS server. The 12d represent the 3D structure of CBLB502 and their backbone comparison with flagellin in which black squares and circles highlight the difference in backbone.*

### A: Overall quality factor of CBLB502 model by ERRAT2 before loop refinement.

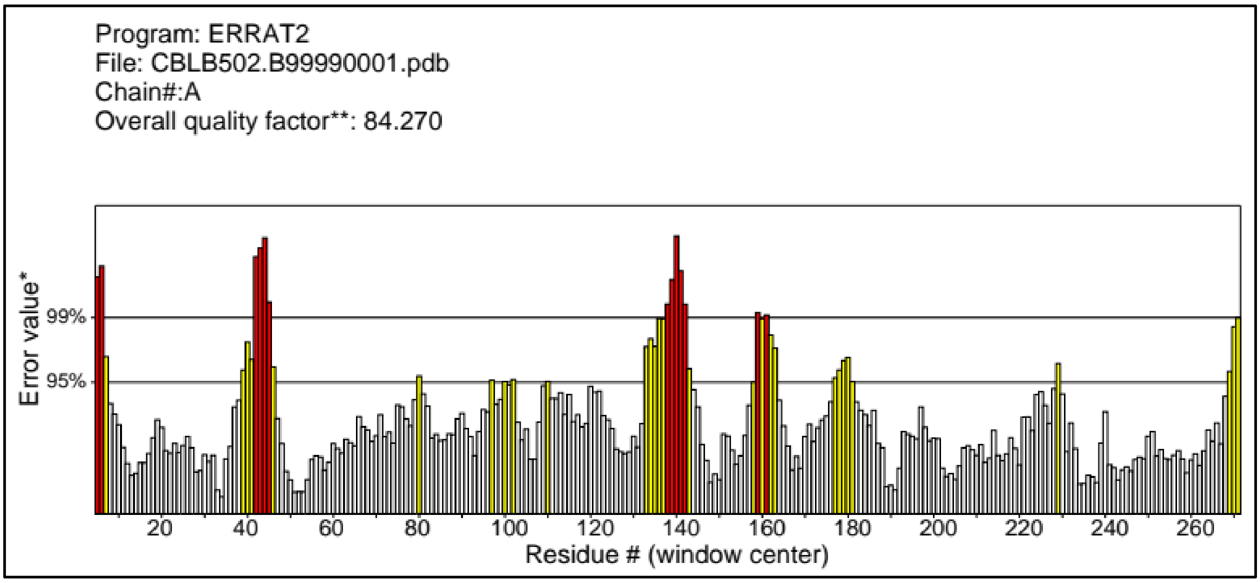


### B: Overall quality factor of CBLB502 model by ERRAT2 after loops refinement

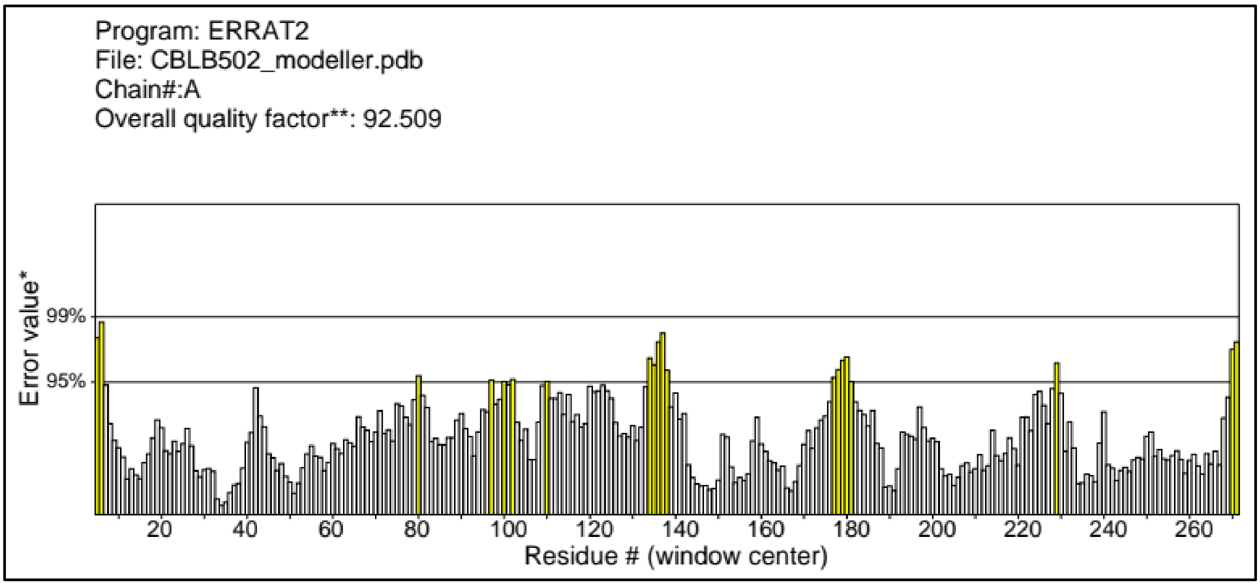


### C: PROCHECK evaluation of CBLB502 model

*This figure suggested the distribution of amino acid residues in allowed (93.4 %) and disallowed regions (0.0%).*

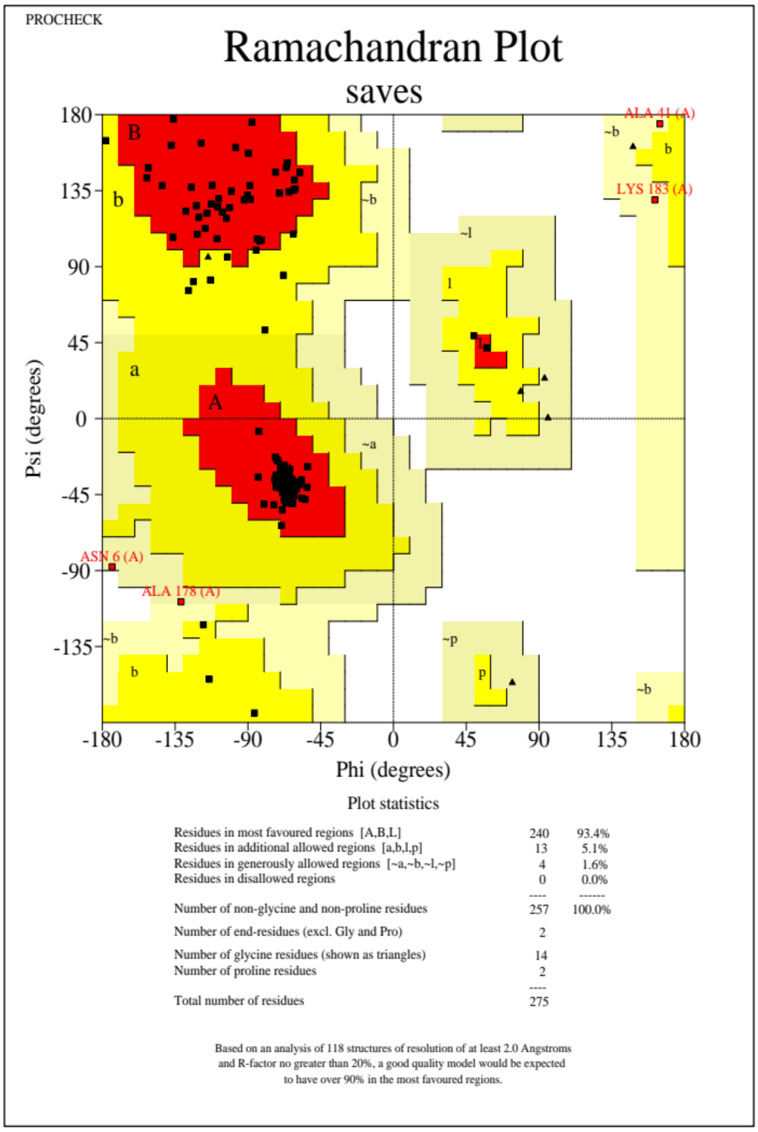


### D: Homology model of refined structure of CBLB502

*The left side of this figure represents the 3D structure of the homology model of CBLB502, having D0 and D1 domains which have similarity with the D1 domain of flagellin on the right side.*

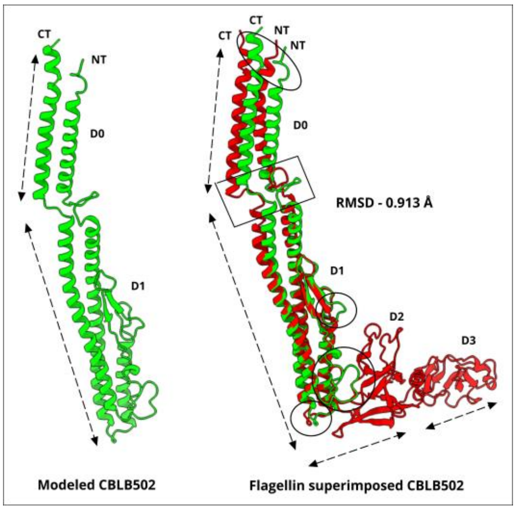

### Supplementary Figure SF4: Intermolecular interactions in TLR5-flagellin and TLR5-CBLB502 complexes.

*This figure highlights the H-bond interactions in both the complexes. The D1 domain of the flagellin (in red) and CBLB502 (in green) are showing to participate in docking with the ectodomain of TLR5 (in blue) and their inset view of intermolecular interactions are represented in the upper and lower side of this figure respectively. We observed more interactions (GLN 98, ASP 114 and ARG 119) in the TLR5-CBLB502 complex than the conserved interactions of both the CBLB502 (ARG91, LEU95, and GLU115) and flagellin (ARG90, LEU94, and GLU114) with TLR5.*

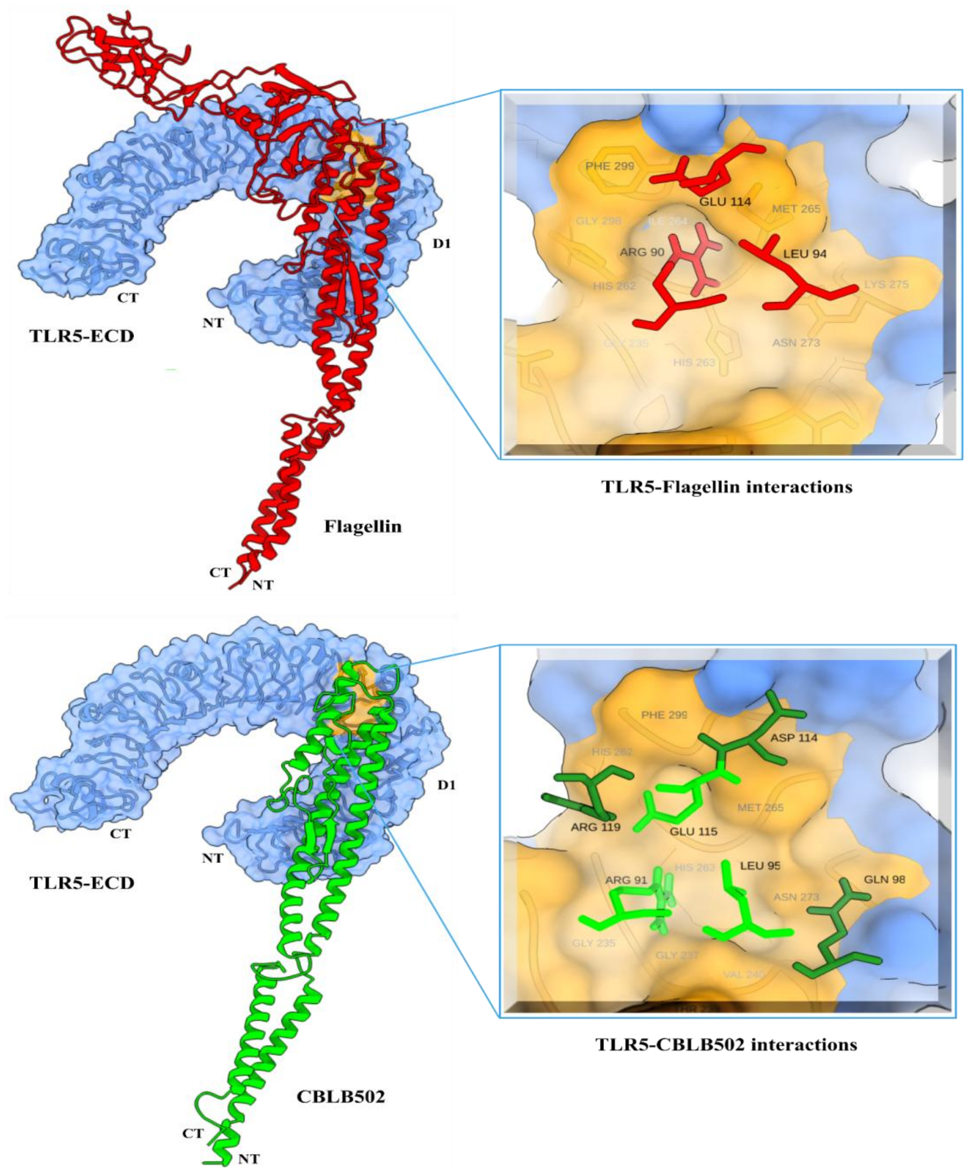

**Supplementary Figure SF5: Impact of CMap-generated leads to other-targets of TLR5 pathway.**

*(A) Correlation of target TLR5 with other proteins of pathway based on TLR5-derived binding affinities of CMap-generated leads. (B) Illustrating grouping of TLR5 pathway proteins including target and other targets based on binding affinities of CMap-generated lead.*

(A)

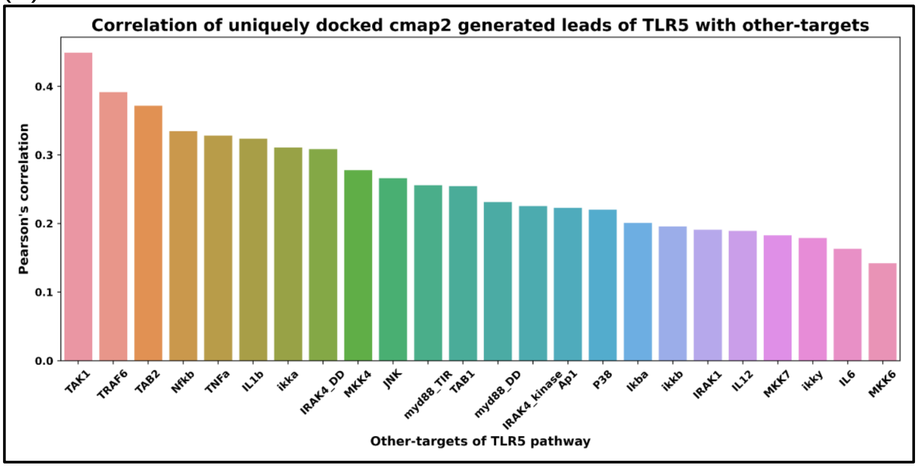


(B)

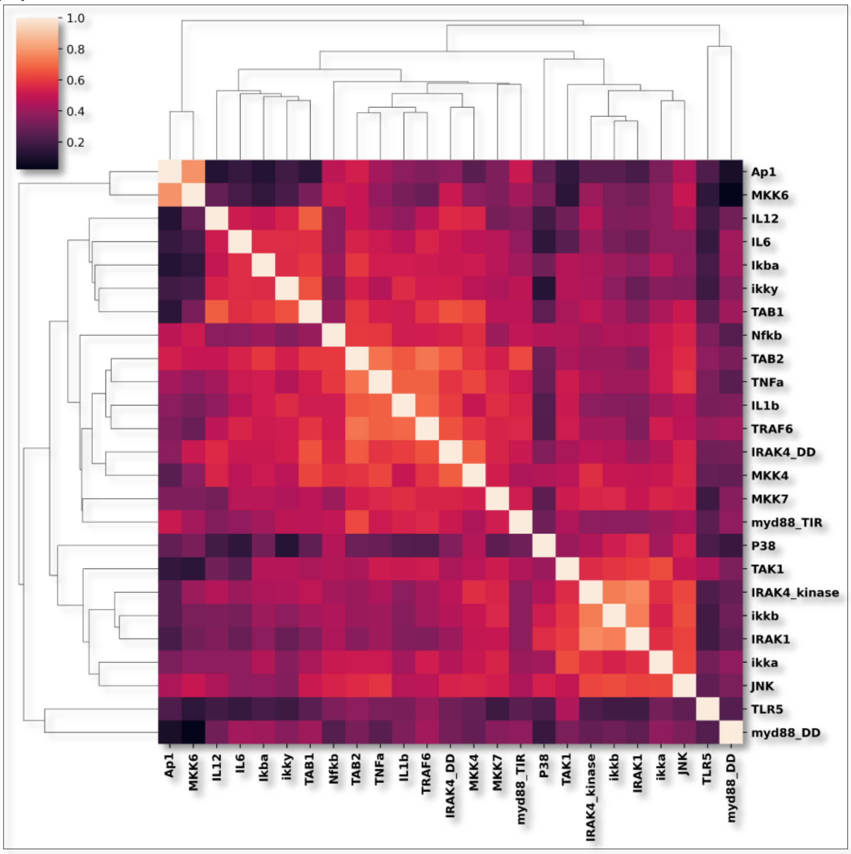
